# Dynamical informational structures characterize the different human brain states of wakefulness and deep sleep

**DOI:** 10.1101/846667

**Authors:** J. A. Galadí, S. Silva Pereira, Y. S. Perl, M.L. Kringelbach, I. Gayte, H. Laufs, E. Tagliazucchi, J. A. Langa, G. Deco

## Abstract

The dynamical activity of the human brain describes an extremely complex energy landscape changing over time and its characterisation is central unsolved problem in neuroscience. We propose a novel mathematical formalism for characterizing how the landscape of attractors sustained by a dynamical system evolves in time. This mathematical formalism is used to distinguish quantitatively and rigorously between the different human brain states of wakefulness and deep sleep. In particular, by using a whole-brain dynamical ansatz integrating the underlying anatomical structure with the local node dynamics based on a Lotka-Volterra description, we compute analytically the *global attractors* of this cooperative system and their associated directed graphs, here called the *informational structures*. The informational structure of the global attractor of a dynamical system describes precisely the past and future behaviour in terms of a directed graph composed of invariant sets (nodes) and their corresponding connections (links). We characterize a brain state by the time variability of these informational structures. This theoretical framework is potentially highly relevant for developing reliable biomarkers of patients with e.g. neuropsychiatric disorders or different levels of coma.

## Introduction

Over the last couple of years, there has been an increasing interest in trying to identify the necessary and sufficient properties of different brain states^1, 2^. One particularly popular theory, the Integrated Information Theory (IIT)^3–7^ proposes a potential route into identifying the essential properties of brain states using five axioms: intrinsic existence, composition, information, integration and exclusion. In other words, given the mechanisms underlying a particular brain state, IIT identifies a brain state with a conceptual structure: an *Informational Object* which is composed of identifiable parts, informative, integrated and maximally irreducible. IIT is thus linked to both *Information Theory* and *Theory of Causality*^8, 9^ but its implementation and applicability to empirical data have remained elusive.

Developing such a framework would be a major step forward, potentially leading not only to a deeper understanding of how the brain process information in different brain states but also to create sensitive and specific biomarkers allowing for the precise assessment of the *Informational Structure* characterising the brain states of individuals in wakefulness, deep sleep, anesthesia, or different levels of coma^2, 10–13^. Yet, thus far, there has been precious little progress in developing the necessary mathematical tools.

Here, we propose an innovative approach to distinguish between different human brain states as measured with neuroimaging. This approach uses an *Informational Structure*; a mathematical structure, related to IIT, which is – as suggested by its name – essentially *informational* and can be used to determine the level of integrated information for different states^14, 15^. *Informational Structure* describes a *global attractor* (GA), a concept from the *Theory of Dynamical Systems*, which determines the asymptotic behavior of a *dynamical system* (DS), i.e. describing the past and future behaviour of the system. GA is a flow-invariant object of the phase space described by a set of selected invariant global solutions of the associated dynamical system, such as stationary points (equilibria), connecting orbits among them, periodic solutions and limit cycles. The informational structure of the GA is defined as a directed graph composed of nodes associated with those invariants and links establishing their connections (see *Methods* and *Supplementary Information* for a formal rigorous definition). *Informational Structure* has been used to show the dependence between the topology, the value of the parameters and the state with respect to its level of integration^14^, as such pointing for the small world configuration of the brain^16^ (although see recent controversies^17^).

Previous research has applied *Informational Structure* to population dynamics in complex networks^18, 19^. Equally used for modelling mutualistic systems in Theoretical Ecology and Economy^20, 21^, *Informational Structure* can relate the topology of complex networks and their dynamics. In mutualistic systems when achieving robustness and life abundance – the so-called architecture of biodiversity^22, 23^ – the dynamics is determined not only by the topology^24^ but also by modularity^25^ and the strength of the parameters^26^.

These kinds of relationships are also at the heart of many important open questions in neuroscience^27, 28^. *Informational Structures* and their continuous time-dependence on the strength of connections can allow to understand the dynamics of the system as a coherent process, whose information is structured, and potentially providing new insights into sudden bifurcations^29^.

Here, we were interested in characterising human sleep, which is traditionally subdivided into different stages that alternate in the course of the night^30, 31^, mainly non-rapid-eye-movement (NREM) and rapid-eye-movement (REM) sleep. NREM is further subdivided into light asleep (N1), NREM sleep stage N2 and deep sleep (NREM sleep stage N3). From N1 to N3, traveling brain waves become slower and more synchronized. In this study, we aim 1) to associate brain states with the variability of *Informational Structure* of GA across time and 2) to demonstrate that this mathematical formalism can be applied to distinguish quantitatively and rigorously between the different brain states of wakefulness and deep sleep using state-of-the-art neuroimaging data collected from healthy human participants.

Dynamical systems can be used to describe a whole range of dynamics; from trivial dynamics, as global convergence to a stationary point, to much more richer ones, usually referred as chaotic dynamics^32–34^ and dynamics of catastrophes^35^. While simple dynamics can be described with total generality, attractors with chaotic dynamics can be only described in details in low dimensional systems. The human brain is a high dimensional system that does not converge or stabilize around a fixed set of invariants. Here, instead they are described as a continuous flow of different GAs and a corresponding flow of *informational structures*. This variability of the *Informational Structure* is assessed assuming a time-varying growth rate parameter of the LV ansatz, which yields a time-varying GA. Moreover, the variability is analyzed applying the Lotka-Volterra Transform, a mathematical operator defined to exactly reproduce the empirical BOLD fMRI signals by finding the growth rate function that tracks their evolution over time. This innovative view of time-varying Informational Structures relies on the fact that we are computing an asymptotic attractor at every time instant, i.e., the GA the system would achieve in the limit assuming a particular value of the growth rate parameter. This is a more dynamical view of the problem which let us adapt to the rich behaviour of the brain. Precisely by characterizing the variability in time of the *Informational Structures* by means of the number of energy levels (see *Models* and *Methods*), we are able to identify the different brain states.

### The Informational Structure (IS) of a dynamical system

This section introduces a mapping between dynamical systems (DSs) and a graph describing its asymptotic solutions and the possible transitions between them.

Usually a DS is described by ordinary or partial differential equations-continuous time-^36^ or difference equations -discrete time-^37^. Generally the phase space *X* of the system comprises the *N* values of the dynamical variables, and is equipped with a family of operators indexed by time, *S*(*t*), defined as *S*(*t*)*u*(*t*_0_) = *u*(*t*_0_ + *t*) for *u* ∈ *X*. A given set 𝒜 is invariant under *S*(*t*) if *S*(*t*) 𝒜 = 𝒜 for every *t*.

We identify the global attractor *𝒢𝒜* with a compact invariant subset of the phase space with the property that *S*(*t*)*u* is arbitrarily close to *𝒢𝒜* for any *u* under the condition that *t* is sufficiently large^38–43^. The structure of *𝒢𝒜* can be given in terms of disjoint subsets and the trajectories in phase space connecting them^44–47^. Given two subsets *A* and *B* of *𝒢𝒜*, they are linked if exists *u* ∈ *X* such that *S*(*t*)*u* is arbitrarily close to *A* for large negative *t* and arbitrarily close to *B* for large positive *t*.

It can be shown that if the DS admits a Lyapunov function^48^, then all the invariant subsets of *𝒢𝒜* can be ordered by connections related to its level of attraction or stability^49^. Thus, *𝒢𝒜* is identified with a directed graph, so that if *𝒢𝒜* = {Ξ_1_,…, Ξ_*n*_}with Ξ_*i*_ disjoint invariant subsets (typically, stationary points or periodic orbits^45, 50–52^, but it could also contain invariant sets with chaotic dynamics^32, 33, 53^), then each Ξ_*i*_ corresponds to a vertex and a directed connection between Ξ_*i*_ and Ξ_*j*_ exists if and only if they are linked by solutions of the DS, in the sense described in the paragraph above. The resulting directed graph is called the Informational Structure (IS), a network describing all the possible future evolution^14, 15, 18, 19^.

These concepts can be illustrated using the Lotka-Volterra (LV) system:

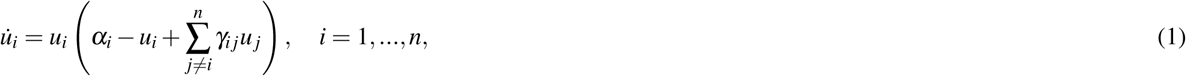

an example of DS on a network (^54–56^) where conditions for the existence and uniqueness of global solutions are well-known^57, 58^.

As shown in Fig.1 with *n* = 2 and suitable coupling matrix *γ*_*i j*_ and *α*_*i*_, there are four stationary points (i.e. invariant subsets comprising only one points in this case). The points are located at the origin (*u*_1_ = 0, *u*_2_ = 0), the x-axis (*u*_1_ ≠ 0, *u*_2_ = 0), the y-axis (*u*_1_ = 0, *u*_2_ ≠ 0), and the positive quadrant (*u*_1_ ≠0, *u*_2_ ≠ 0). Thus, in this case the *𝒢𝒜* is comprised of four points in ℝ^2^. Each stationary point is hyperbolic and locally creates a field of directions towards (stability) or from them (instabilities). The stationary points of the LV system for *n* = 2 can be described as two binary variables, null values indicating that the corresponding dynamical variable has zero value (i.e. is located in one of the axes), and non-null values indicating that it is located outside that axis, e.g the stationary points can be label as (0, 0), (0, 1), (1, 0), and (1, 1). In other words, the *𝒢𝒜* of the system (and thus its asymptotic behavior) could be encoded as a set of Boolean variables and connecting global solutions among them.

**Figure 1.**
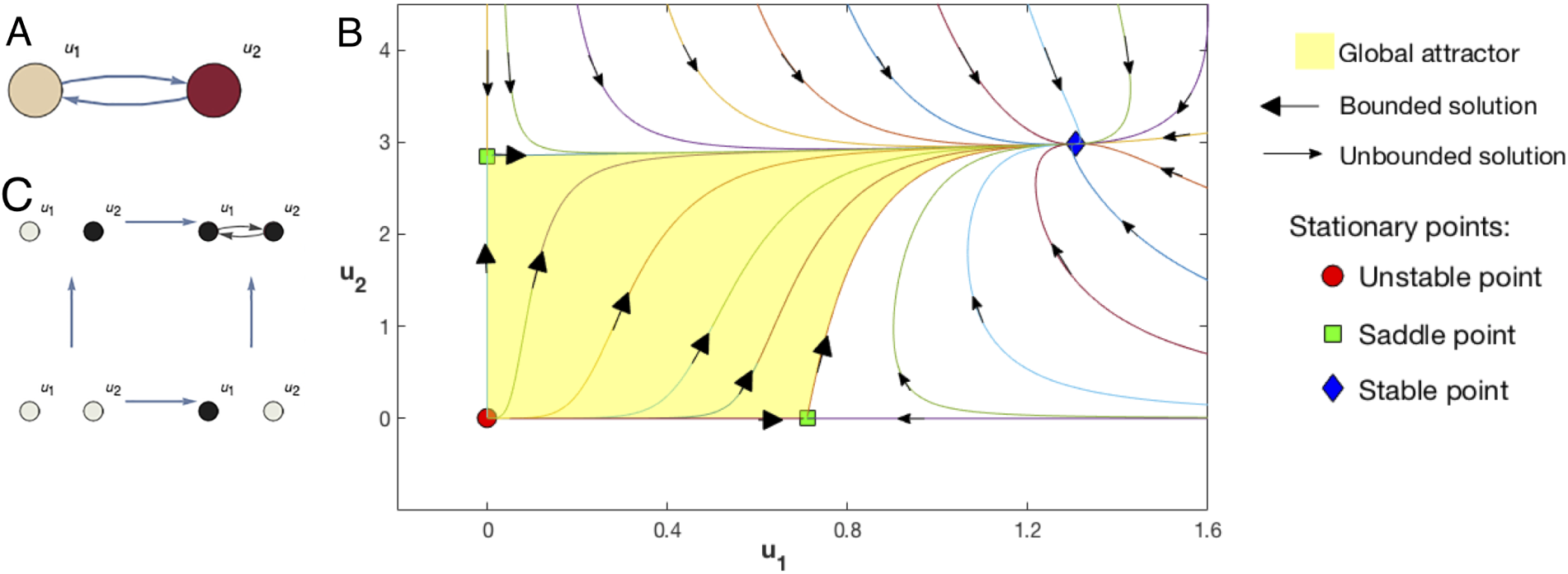
**A**, Structural network of a 2-dimensional (*n* = 2) Lotka-Volterra (LV) system given by (1) (see Supplementary Information for *α* and *gγ* values in this example). **B**, Solutions of that LV system in the phase space. The global attractor is the set of all bounded solutions. There are one trivial and unstable solution (red circle) and two saddle points (green squares) both semitrivial stationary points, one in the *u*_1_ axis and another in the *u*_2_ axis. Each saddle point is hyperbolic and locally creates a field of directions towards (stability) or from them (instabilities). The stable point (blue diamond) is a stationary point with two strictly positive components.**C**, The Informational Structure (IS) is a new network made by four nodes associated with the four stationary points and directed links associated to bounded solutions. The relation induced by the links is transitive in such a way that only the minimal links to understand the connectivity between nodes are represented. Each node of the IS can be represented as a subgraph of the original LV system where non-null components of the associated stationary point are shown in black and null in grey. In this example there are 3 energy levels: the trivial solution, the saddle points, and the stable solution.

The example based on the LV system for *n* = 2 in Fig.1 shows how the stationary points are partially ordered, i.e. they admit an order relationship in which not necessarily all the pairs of elements can be compared. In general, two invariant subsets Ξ_*i*_ and Ξ_*j*_ can be compared if and only if there exists a chain of invariant subsets Ξ_*l*_, *i* ≤ *l* ≤ *j* that are linked in the phase space, and are ordered according to their level of attraction or stability, as given by the Lyapunov function^49^ (see also^59^). In the case of the LV equations for *n* = 2, the unstable and stable points are maximal and minimal in this ordering, respectively, with the other two stationary points not being ordered one with respect to the other.

It can be shown that the energy levels *𝒩* = *N*_1_, *N*_2_,…….., *N*_*q*_, are well defined and indicate the level of attraction or stability. The number of energy levels (NoEL) *q* will be the most important parameter in our data analysis (see Supplementary Information including Fig. 2.2 for more details and formal definitions).

### Measuring energy levels from fMRI data

The autonomous LV equations have been used to generate reproducible transient sequences in neural circuits^60–64^, but global brain dynamics do not converge or stabilize around a fixed set of invariants, and might be described as a continuous flow of quick and irregular oscillations^27, 65^. Thus, instead of describing dynamics in terms of asymptotic behavior, an alternative consist of introducing the dynamical informational structures (DISs), defined as the ISs indexed continuously by time (see Fig.2).

**Figure 2.**
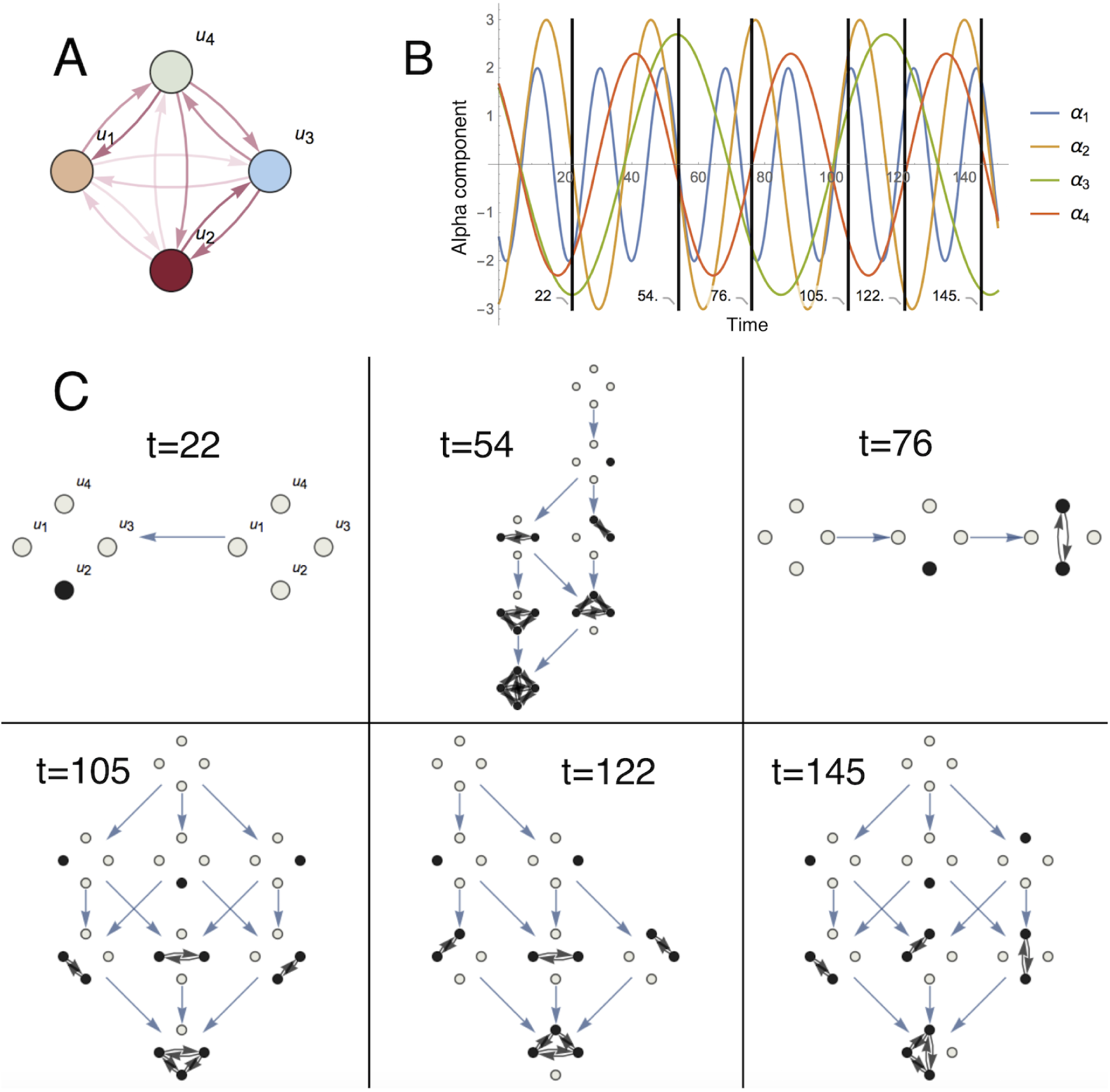
**A**, Structural network of a 4-dimensional (*n* = 4) Lotka-Volterra (LV) system given by (1) (see Supplementary Information for *gγ* values in this example). **B**, Evolution in time of *α*_*i*_ parameters of a four-node system. In this example *α*_*i*_ are periodic functions of time. Changes in the parameters governing the dynamics produce changes in the corresponding informational structures (ISs). **C**, Dynamic Informational Structures: different ISs corresponding to the time steps shown above. For the cooperative LV system each energy level is formed by nodes associated with stationary points with the same number of non-zero components. The number of energy levels (NoEL) *q* changes over time also, but equals the number of non-zero entries in the GASS plus one (see Supplementary Information for more details about the NoEL).

We consider experimental data from a DS whose underlying generating differential equations are unknown (in this case, fMRI data). We can select the parameters in the LV system so that the data satisfies a non-autonomous version of the equations. In other words, we can select the parameters from time to time, so that the dynamics of the system evolves exactly according to the LV equations. Then, these equations can be used to identify the DISs of the system, consisting of the temporal evolution of the partial order between Boolean variables that encodes the stationary points and the solutions connecting them. Thus, the LV equations are not used to model the temporal evolution of the system itself, but as an ansatz to obtain the temporal evolution of the *𝒢𝒜* s and associated DISs.

For our purpose, we consider the non-autonomous LV equations, given by:

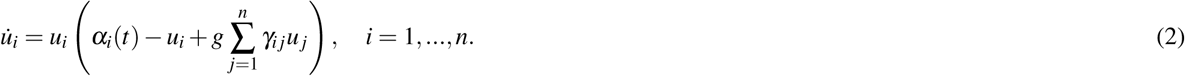

From these equations we can obtain the time-dependent parameters *α*_*i*_(*t*) as:

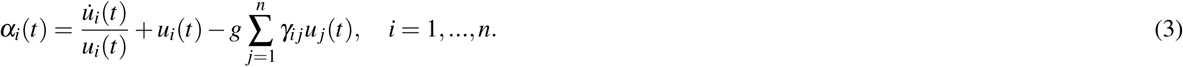

This equation defines the Lotka-Volterra transform (LVT), i.e. a non-autonomous LV system that fits exactly the empirical data and its first derivative. In practice, both *u*(*t*) and 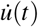 will be empirical values from BOLD signals obtained using fMRI in the form of discrete time series.

It can be shown that an equivalence exists between obtaining the stationary solutions of LV systems and solving a linear complementarity problem (see^58, 66, 67^). For Lyapunov stable systems^68, 69^, any initial point of the interior of the positive quadrant will converge to the globally asymptotically stable solution (GASS). Equivalently, the IS associated with the *G A* is a directed graph such that a single stationary GASS is located at the minimum energy level. Furthermore, the number of energy levels (NoEL) *q* equals the number of non-zero entries in the GASS plus one (see Fig.2). The GASS is associated with a node or vertex with only incoming edges, implying stability. The linear complementarity problem associated with identifying the *α*_*i*_(*t*) can be solved using the Complementary Pivot Algorithm^66, 67^.

In our LV equations *n* = 90 and for each time there are up to 2^90^ = 1.238 × 10^27^ stationary points, each one a unique combination of binary variables. We transform the time series of ISs into time series of NoEL, and investigate their distributions during wakefulness and deep sleep (see Supplementary Information for more details and formal definitions).

### Identifying brain states: paired difference test and J index

To assess differences in the DISs between wakefulness and deep sleep, we use the non-parametric Wilcoxon signed-rank test. We consider two quantitative descriptors of the DISs, the time average of NoEL 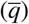 and the standard deviation of NoEL (*σ*_*q*_), both calculated for each participant in the two brain states. We obtained the p-value of a paired and two-sided test for the null hypothesis that the distribution of average NoEL (or their standard deviation) presented the same median during awake vs. deep asleep subjects.

In addition, we defined the following measure

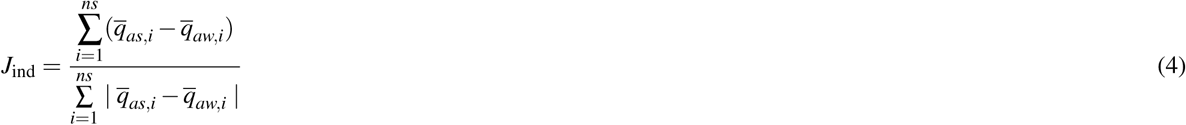

where *ns* is the total number of subjects, 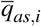 is the value of 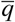 for subject *i* while asleep and 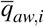 is the value of 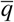 for subject *i* while awake. Then, *J*_ind_ = 1 (*J*_ind_ = −1) if the mean number of energy levels is larger (smaller) for deep sleep than wakefulness for all subjects and *J*_ind_ ≃ 0 if the null hypothesis holds. Please note that *J*_ind_ can also be defined for *σ*_*q*_ or for any other parameter (see Supplementary Information for more details).

## Results

We aimed characterize the different brain states of wakefulness and deep sleep by analyzing the variability of the Informational Structure (IS) across time as shown in Fig.3 with the flowchart of steps. Briefly, first we created a whole-brain ansatz based on the Lotka-Volterra (LV) equations for the brain dynamics, where the structural connectivity matrix obtained from empirical tractography-based dMRI is chosen as the interaction matrix of the dynamical system (DS). Second, we applied the Lotka-Volterra Transform (LVT) to calculate the growth rate *α* that reproduces exactly the filtered empirical BOLD fMRI signals. Third, after solving the linear complementarity problem we computed the globally asymptotically stable solution (GASS) and the number of energy levels (NoEL) of the corresponding IS at each time instant (Fig.3). Finally, in order to assess the differences between the distributions of the average NoEL 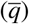 and decide which brain state the participant is in, we applied statistical tests, namely Wilcoxon and *J*_ind_ as defined in (4).

**Figure 3.**
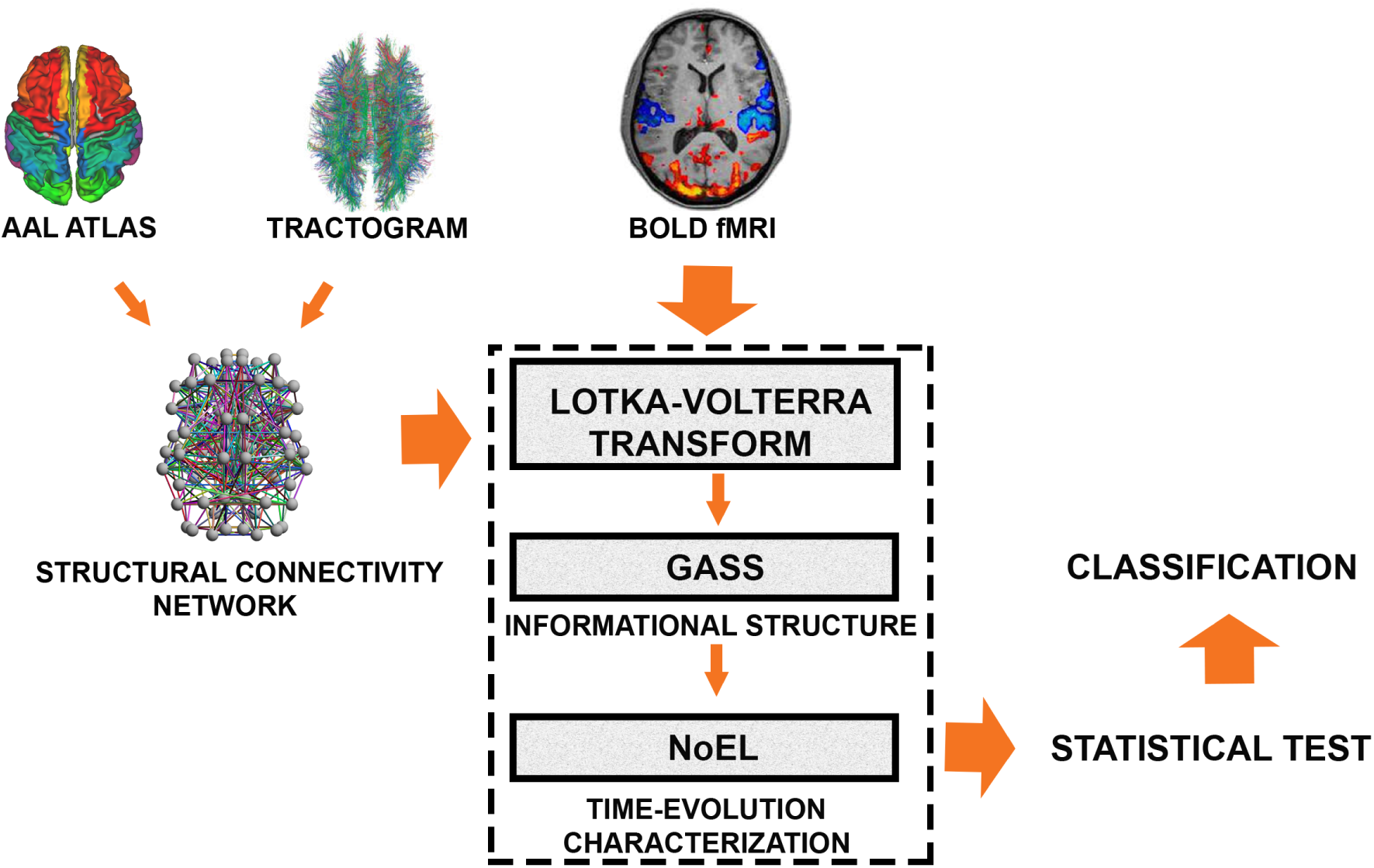
Flowchart illustrating the methods. For this study, we used the structural connectivity between 90 standardized brain areas through an automated anatomical labelling (AAL) atlas. The corresponding tractography was obtained in a previous study^70^ through diffusion magnetic resonance imaging (dMRI) providing an structural connectivity network. We assume that neural activity in different brain areas can be simulated in the ansatz of interacting species in a cooperative Lotka-Volterra (LV) system constrained by the structural connectivity and a global coupling strength parameter value. With these constraints, the ansatz defines the LV transform, a mathematical operator that calculates the growth rate as a time function that reproduces exactly the filtered empirical BOLD fMRI signals. Then solving the linear complementarity problem we calculate the globally asymptotically stable stationary solution (GASS) and the number of energy levels (NoEL) of the corresponding informational structure at each time instant. In order to assess how different the distributions of the NoEL for awake and for deep asleep are, we calculate a statistical hypothesis test (Wilcoxon test) and the *J*_ind_ (see Fig. 2.3 in Supplementary Information for a more detailed flowchart).

### Lotka-Volterra ansatz fitting

#### Impact of coupling strength

The strength of the connections between pairs of nodes is controlled by the coupling strength parameter. Changes in coupling strength *g* led to changes in *α*, and therefore had an impact on the distribution of NoEL. Fig.A shows the impact of *g* on the ability of IS to differentiate between the two brain states, i.e. demonstrating how the gap between awake and deep asleep measures changes as a function of *g*. A vertical dashed line showing the maximum value (*g* < 0.36743) ensuring global stability is shown. We computed the Pearson linear correlation coefficient between the two curves (−0.9781) to corroborate that the *J*_ind_ basically measures the same as the Wilcoxon test (although *J*_ind_ is more intuitive and easier to calculate). The optimal value (*g* = 0.29) was the same in both tests, and we chose it for our computations. Note however that the gap between awake and asleep was not restricted to this value. Indeed, the range of *g* in which the differences between the distributions of energy are manifest (p-values below 1% or 10^−2^) was very broad. Fig.A shows that there was a wide range of values for which the p-value was small: from *g* = 0.05 to the red dashed line where the global stability of the system is not guaranteed. In that same interval *J*_ind_ remained above 72% (*J*_ind_ = 0.72). In other words, our results were robust, as they do not depend on a specific value of *g*. In addition we verified that the suggested method does not produce good results when *g* tends to zero or when *g* approaches the maximum limit guaranteeing stability of the system.

#### Impact of filtering

Fig.B shows the results of the Wilcoxon test and *J*_ind_ index for the means 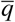 and the standard deviations *σ*_*q*_ of the NoEL assuming different filter ranges. At high frequencies, noise was not filtered and the DISs of the two brain states were very similar, the p-value was high and the *J*_ind_ small. For 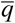 the filter range with minimum p-value was 0.077−0.096 Hz (*p* = 2.3323·10^−4^). For *σ*_*q*_, the filter range with minimum p-value was 0.077−0.106 Hz (*p* = 2.3323·10^−4^) and the second best was 0.077−0.096 Hz (*p*-value= 2.7642·10^−4^). For *σ*_*q*_, the filter range with maximum *J*_ind_ was 0.077−0.106 Hz (*J*_ind_ = 0.9954) and the second best was 0.077−0.096 Hz (*J*_ind_ = 0.9813). Therefore, we chose the filtering range 0.077−0.096 Hz. When the signal was filtered in this range the difference between the distributions of NoEL and, therefore, between the DISs of awake and sleeping participants was maximal. The values obtained for most filters (wide filters, narrow filters or even filter absence, i.e. 0.0−0.24 Hz where 0.24 Hz is the Nyquist frequency) were good enough (*J*_ind_ > 0.75 and *p*-value< 0.01) to sustain that our results can not be attributed to a specific filter. Rather, we demonstrated that our hypotheses, in general, were true even for the unfiltered signal. Only high-pass filters provided negative results (*J*_ind_ < 0.4 and *p* > 0.2). Thus, the results did not depend on a specific filter and our conclusions were again robust in this regard.

#### Statistical analysis of IS variability

The Lotka-Volterra Transform (LVT) allowed us to compute the time-varying vector *α* for every participant in both brain states. Using the LCP we computed the globally asymptotically stable solution (GASS).

Fig.5 shows a 60-seconds long sample of the NoEL as a function of time for one subject in awake state and deep sleep state. The large and the small number of energy levels were more frequent in awake than in deep asleep conditions. Furthermore the average NoEL appeared to be higher for deep sleep state than for the awake state. To confirm the latter we studied the NoEL distributions for the 18 participants in both conditions. Generally, the NoEL increased when the components of alpha grow (Fig.6A), where alpha components were computed using the LVT. Fig.6B shows the distribution of energy for every participants in awake and in deep sleep conditions back to back. We see that for both conditions the value *q* = 91 was the most frequent one, whereas *q* = 90 was the second most frequent. When *q* = 91 the corresponding ISs had the maximum NoEL (the GASS has all the 90 components different from zero), i.e., the attractor (the GASS) was comprised by the whole brain (all brain areas were active). This whole brain attractor, which was more frequent in awake than in deep asleep conditions, was a consequence of hubs simultaneously increasing their activity. Generally, NoEL distributions for awake were very different from those for deep sleep phase in every participant.

**Figure 4.**
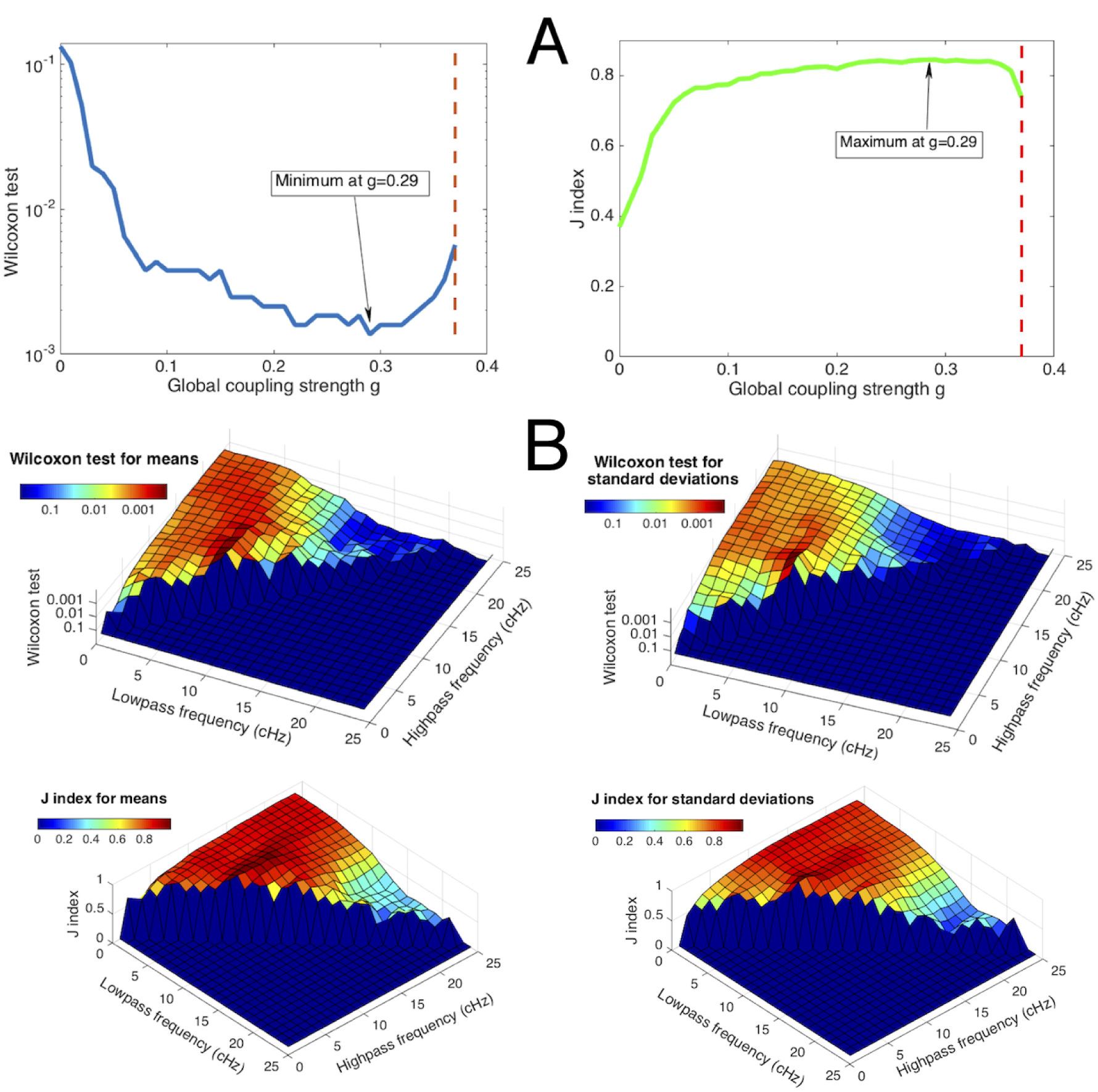
**A**, Fitting of the coupling strength parameter *g* taking as an optimization criterion the minimization of the p-value of the Wilcoxon test and the *J*_ind_ maximization, both for means, as optimization criteria. The red dashed line delimits the region in which global stability is guaranteed. In both cases the optimal value is *g* = 0.29. **B**, Fitting of the filter used taking as an optimization criterion the minimization of the p-value of the Wilcoxon test and the *J*_ind_ for the samples of both, 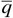 and *σ*_*q*_. Both, the Wilcoxon test p-value and the *J*_ind_ are functions of the ends of the filtering range (see Fig. 3.1 in Supplementary Information for a 2D version of the graphics).

**Figure 5.**
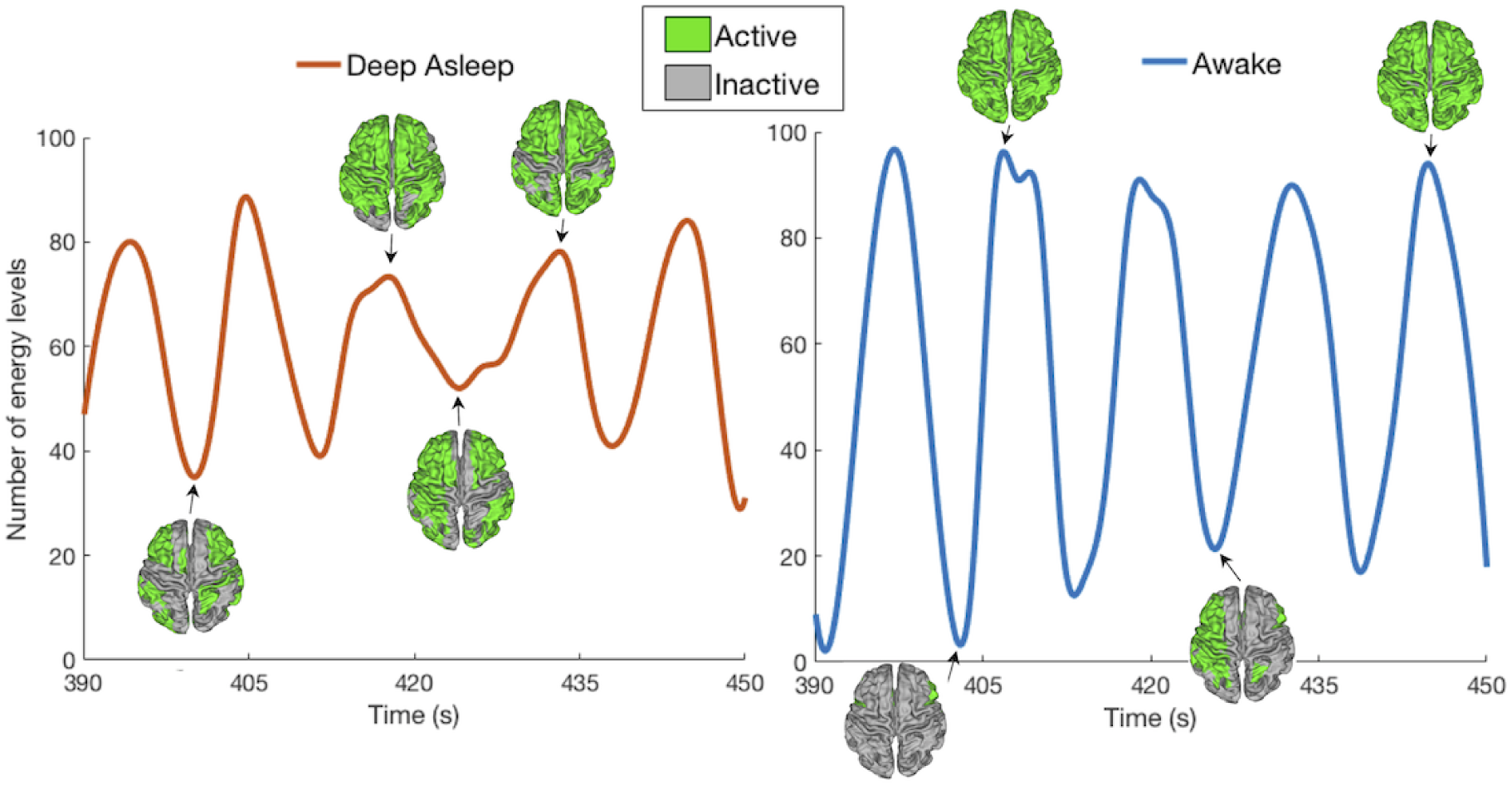
Example of a 60-seconds long sample of the number of energy levels (NoEL) for one participant in deep sleep and wakefulness states. Empirical data was band-pass filtered in the range 0.077−0.096 Hz and *g* = 0.29. In practice both the BOLD signal data and the NoEL were evaluated as time series, that is, for discrete values of time (Δ*t* = 2.08 seconds). Only for this figure the discrete data were interpolated using cubic spline. This sample shows greater variability for awake than for deep asleep condition. Furthermore, the average NoEL was higher for deep asleep than for awake condition (see Fig. 3.2 in Supplementary Information for a similar graphic using a different filter).

**Figure 6.**
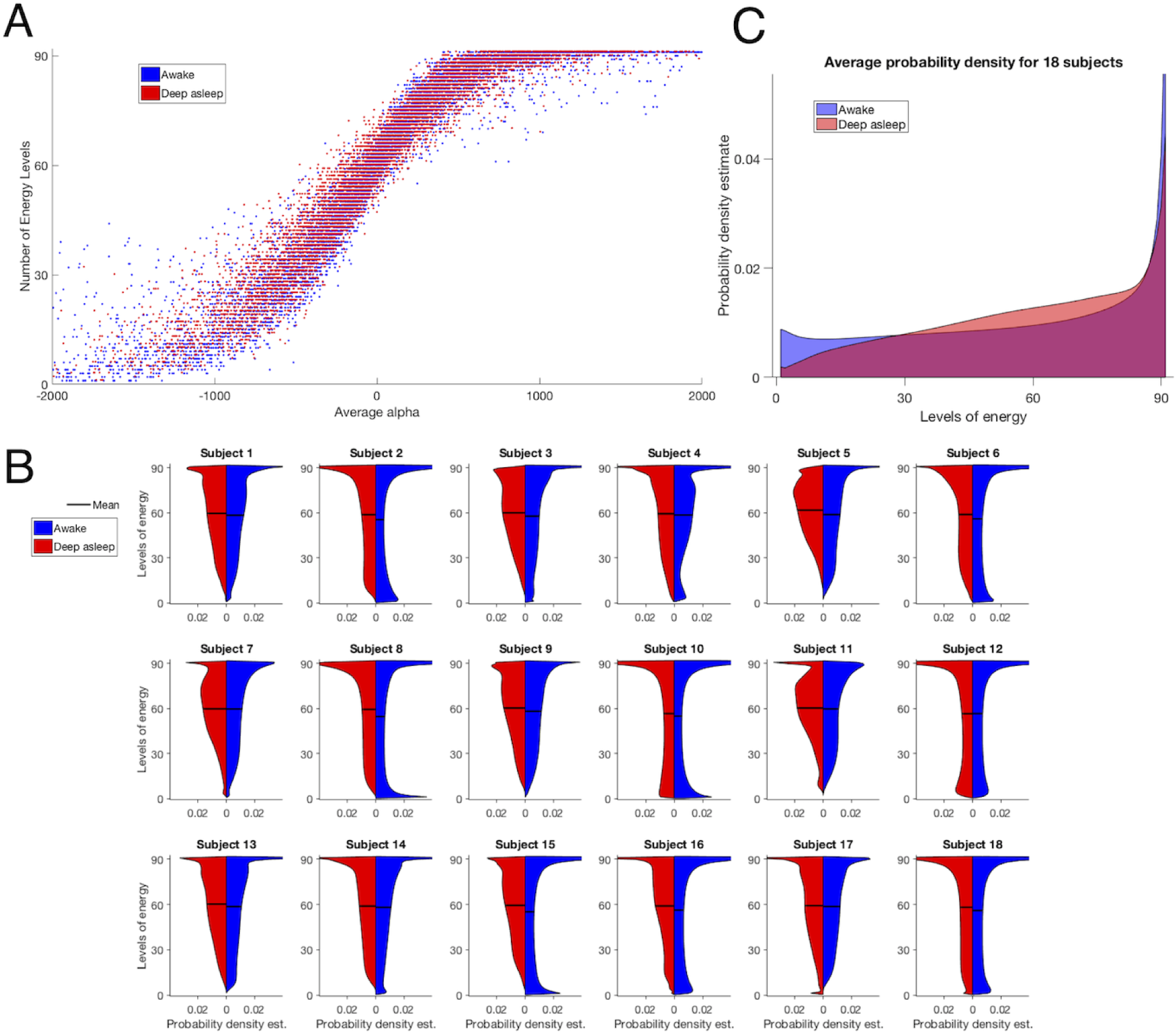
**A**, Number of Energy Levels (NoEL) as a function of the 90 alpha components on average. Each dot represents a time point of a subject in a brain state. **B**,Distribution estimations of the NoEL for 18 healthy participants in awake and in deep sleep states back to back. The smoothing was made using the kernel density estimation, i.e. a non-parametric way to estimate the probability density function. Inferences about the population are made based on a finite data sample (see Fig. 3.3 in the Supplementary Information for the non-smoothed histograms). The NoEL was obtained from filtered data between 0.077− 0.096 Hz. **C**, Average distribution over the 18 participants in wakefulness and deep sleep conditions (see Fig. 3.4 in the Supplementary Information for the non-smoothed histogram and the back to back probability density estimation).

There seemed to be a tendency for extreme values of NoEL to be more frequent when the participant was awake than asleep, while for the medium-high values of NoEL the opposite held. The pattern observed was that awake distributions (blue) were more homogeneously distributed among all the possible values of *q*, whereas asleep distributions (red) tended to have higher values (> 45) of NoEL. In general, this pattern was repeated in most of the individuals, however there was a striking inter-individual variability. Please note that for participant 12 the tendency was reversed and the distributions for the two brain states were almost symmetric. Apart from this case, the distributions of energy in awake and in sleep states were clearly different for all other participants.

In order to draw general conclusions beyond the inter-individual differences, we computed the average distribution of the 18 participants in the awake state and the average distribution of the 18 participants in the sleep state (Fig.6C). Although in all cases the highest NoEL was frequent, the most interesting patterns were observed when the frequencies for the maximum values *q* ∈ [80, 91], were not taken into account. When looking at the frequencies below 80, the probability grew linearly with the NoEL for the deep sleep condition, whereas it showed almost a constant behaviour for the wakefulness condition with *q* ∈ [1, 80], as it can be seen from Fig.6C (red and blue respectively). The pattern typically observed was that the distribution for the awake state was more homogeneous along all the possible NoEL values. ISs with extreme NoEL values, that is, low (< 28) or very high (90 or 91), were more frequent in awake individuals as suggested by the sample of Fig.5.

In Fig.7A the error bars were computed according to 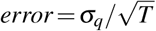. Therefore, in those individuals in which the time series were shorter (shorter time spent in deep sleep or wakefulness) the error bars are overlapping. With filtering in the range 0.077− 0.096 Hz for all participants except participant 12 the mean NoEL 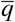, was higher when the participants were in deep sleep. In order to assess how different 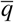 was for participants in awake and deep sleep, we used the signed-rank test Wilcoxon (see section 2.5) and we obtained *p* = 2.3323·10^−4^. In addition, we computed the *J*_ind_ index defined in (4) and obtained *J*_ind_ = 0.9948. Recall that *J*_ind_ = 1 when 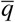 was larger for all participants in the deep sleep state. Please note that there are up to six participants for whom the two error bars did not overlap.

**Figure 7.**
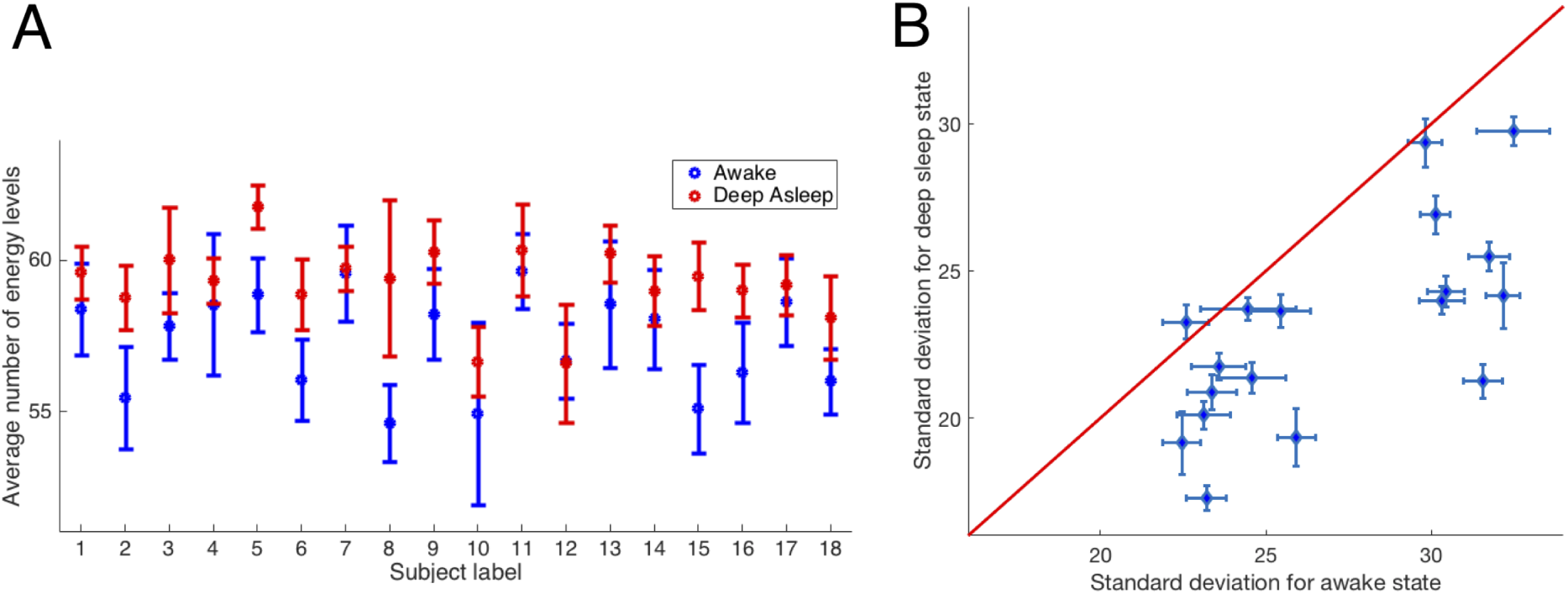
**A**, Average number of energy levels (average NoEL) 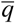 for a Lotka-Volterra ansatz with *α*(*t*) fitted to empirical data of 18 participants awake (blue) and deep asleep (red) (*J*_ind_ = 0.9948, *p* = 2.3323·10^−4^). Differences between the means of both brain states were significant. The error bars were calculated according to 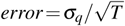 where *σ*_*q*_ is the standard deviation of the NoEL and *T* was the length of the data series. **B**, Standard deviation of the NoEL for deep asleep state as a function of NoEL standard deviation for wakefulness state. The standard deviation was the same for wakefulness and for deep sleep brain states along the red line. Clearly, in 17 out of 18 participants the standard deviation was larger in awake than in deep sleep brain state (*J*_ind_ =−0.9813, p-value= 2.7642·10^−4^). Again, the differences between the standard deviations of both brain states were very significant (see Fig. 3.5 in the Supplementary Information for a similar graphic using other filters).

Then, we established patterns regarding the variability of the distributions. We computed the standard deviations of the distribution of NoEL in each participant for the two brain states. In Fig.7B each point represents a participant. For 17 (out of 18) participants the standard deviation was larger for wakefulness than for the deep sleep state, *J*_ind_ =−0.9813 and *p* = 2.7642·10^−4^. Again, the differences between the standard deviations of both brain states were very significant. There was only one participant who shows a greater deviation when asleep. This may be clearly explained recall from Fig.6C that, whereas the standard deviation for the awake conditions showed an almost constant behaviour, with values of *q* homogeneously distributed for all levels of energy, the standard deviation for the deep asleep condition increases almost linearly with the levels of energy, with a greater concentration in the larger values of NoEL.

Next, we studied what were the active areas (non-zero components) in the attractor (GASS). Given that, on average, the attractor of the deep sleep state was more populated (larger NoEL) than that of the wakefulness state. In addition, most of the brain areas tended to appear more frequently in the deep sleep attractor than in the wakefulness attractor. In other words, as 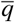 was greater in the participants in deep sleep, the general tendency was that each area of the brain was more frequently present in the deep sleep attractors than in the wakefulness attractors (with some exceptions). As shown in Fig.8AB, we computed the frequency of appearance of an area in the GASS for each participant and for each brain state and then averaged over all participants. We computed the difference between the frequencies in awake and in deep sleep states for each participant and area (Fig.8C). As can be seen, most of the areas showed less presence in the wakefulness attractor than in the deep sleep attractor. But there were some exceptions (indicated by the warm colours), i.e., greater presence of the area in the wakefulness attractor (see Fig. 3.6 in Supplementary information for differences across participants, where again we found striking inter-individual variability but where, in the majority of cases, warm colors were exceptional).

**Figure 8.**
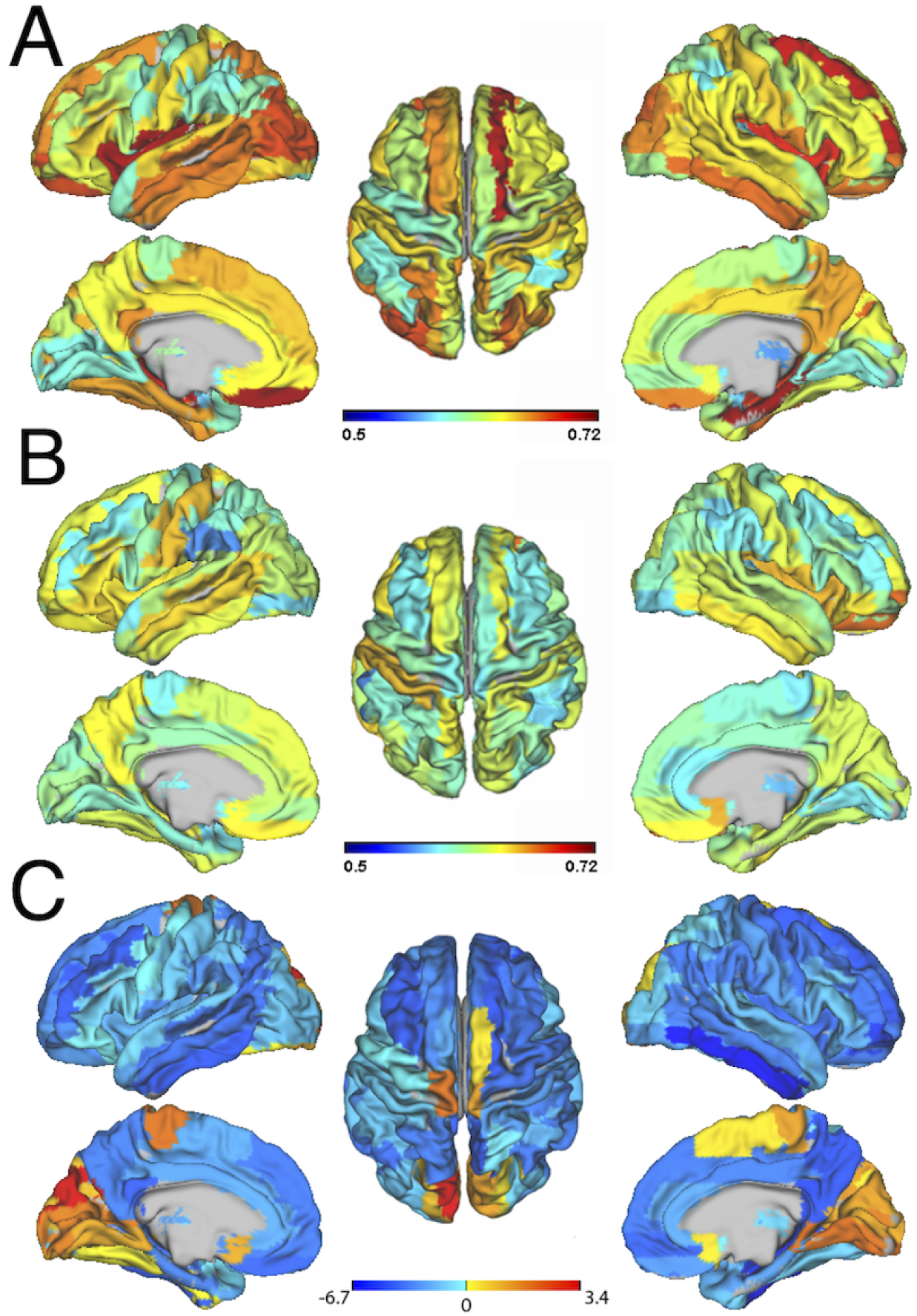
Frequencies of appearance of each area in the globally asymptotically solution (GASS) in deep sleep (**A**) and in the wakefulness (**B**) states averaged over all participants. **C**, Difference between the averaged frequencies of appearance of each area in the GASS in both states. This difference is expressed in standard error units where the standard error is estimated by the interpersonal variability. Most areas have cold colours, i.e., greater presence in the deep sleep attractor (see Fig. 3.6 in the Supplementary Information for more details and individual differences).

Taking into account the averaged results (see Fig.8) for the 18 participants, 14 areas (out of 90) showed this opposite tendency (see Fig.8). 12 out of these 14 are actually six pairs of homotopic zones, i.e., pairs of areas that occupy symmetrical zones in each hemisphere. These six pairs of areas are the olfactory, the calcarine (V1), the cuneus (basic visual), the lingual (visual letters), the occipital superior (visual), and the paracentral lobule (motor-sensory), all of them found in both left and right hemispheres. Furthermore, the fusiform in the left hemisphere (facial recognition) and the supplementary motor area in the right hemisphere (control movement). Due to their specific functions, in all cases, it makes sense that these areas were foci of attraction in the wakefulness state.

## Discussion

Here we have proposed a novel mathematical formalism for characterizing the time-varying attractors landscape depicted by the dynamical activity of the human brain. In this rich and complex landscape, finding the energy levels of the informational structure (IS) of a global attractor *𝒢𝒜* could be seen as identifying the maximum stability in a landscape that changes with time. Our novel method allows us to clearly characterise and distinguish between different brain states, opening up for relevant clinical applications in terms of improving existing biomarkers.

Specifically, the results show that the method is able to significantly characterise and distinguish between the well-defined brain states of wakefulness and deep sleep (N3 stage) in empirical BOLD functional magnetic resonance imaging (fMRI) signals (as measured with concurrent EEG and which helped trained neurologists to identified the sleep stage). For this purpose, we used a whole-brain dynamical *ansatz*. This ansatz integrates the underlying anatomical structure, obtained from tractography-based diffusion magnetic resonance imaging (dMRI) with functional local node dynamics measured with fMRI. In contrast to previous studies which have used such nodes as the regions of interest^71–73^ in a volume-based human brain atlas^74–77^, here we instead used the neural activity at each node assuming a cooperative Lotka-Volterra (LV) ansatz (see Methods). LV is a well-known system of first-order nonlinear differential equations used to describe the dynamics of biological systems in which several species interact^58, 69, 78^. The GA for this model consists of the equilibria and the heteroclinic connections among them, and its characterization is, to some extent, well understood^57, 69^. The informational structure of this GA can be computed and, under some conditions, there exists a finite number of equilibria and directed connections among them that create a hierarchical organization with several levels of equilibria sets or energy levels ordered by connections in a gradient-like fashion^18, 19^.

We assessed the variability in the human brain by analyzing the time-varying IS, the graph associated with the *𝒢𝒜* of a dynamical system (DS). Our approach was based on the Lotka-Volterra (LV) equations for collaborating species in a whole-brain ansatz. The growth rate parameter *α* was assumed time-varying, and a time series of ISs was obtained. We defined the Lotka-Volterra transform (LVT) as the growth rate parameter, *α* = *α*(*t*), that reproduces exactly the empirical BOLD signal. This allowed us to compare the corresponding dynamical informational structures (DISs) in wakefulness and deep sleep conditions.

The first remarkable thing from our results is the inter-individual variability shown by the participants, which is greater than what would be expected beforehand (see Fig.6B). The human brain is characterized by a surprising inter-individual variability in neuroanatomy and function that is reflected in large individual differences in human cognition and behaviour ^79^. This diversity is undoubtedly one of the marks of our species.

Still, we have shown that the ISs can be used for the study of brain dynamics, and to compare and distinguish between different brain states. Indeed, the main conclusion of this study is that are significant differences in the ISs corresponding to the brain activity of individuals in wakefulness and deep sleep. Before seeing the probability density estimates in Fig.6BC, unimodal distributions such as Gaussian distributions could be expected. At least it would be reasonable to expect two similar distributions but with different maxima. However, not only the means of the number of energy levels (NoEL), *q*, are different, but also the distributions themselves. Besides the very high frequencies obtained for the highest values of *q* in both distributions, the frequency of the deep sleep NoELs increases almost linearly with *q*, while that of wakefulness remains almost constant in most *q*.

On the other hand, the ISs with the highest NoEL, and therefore possibly those with the highest number of nodes and vertices, are the most repeated for participants both in wakefulness and deep sleep. In addition, in the awake state the maximum NoEL is more frequent than in sleep condition. Recall that this attractor or globally asymptotically stable solution (GASS) has all the components different from zero, that is, it consists of the active complete brain (the 90 active areas) which implies a high degree of effective connectivity. Therefore our results agree with the fact that during slow-wave sleep effective connectivity decreases ^80^.

Generally, the extreme number of energy levels are more frequent when awake compared to deep sleep as shown in Figs.5, 6 and 7B. These results are an effect on the global level of a group of local nodes with a number of links that greatly exceeds the average (hubs) driving the dynamic system when these nodes simultaneously increase or decrease their activity. It could suggest a higher capability to integrate information whilst awake and a increased capacity to amplify local perturbations. In contrast, slow-wave deep sleep is associated with a diminished level of information integration^81–84^. This result is compatible with the integrated information theory of consciousness^3–7^, which holds that different levels of consciousness must correspond to the capacity of the brain to integrate information. Many other studies have shown that integration is impaired during non-wakefulness^10, 85, 86^.

Our results show also a decrease of time variability in deep sleep condition that could be associated with a decrease in the differentiation of brain activity. According to the information-integration theory this might be thought of as an indicator for diminished conscious awareness^4^.

It is worth mentioning that we have not observed any NoEL or IS that are exclusive of a particular brain state. Any NoEL or IS can be found for any of the two states, and it is only through its time variability that we can asses the differences between the brain states.

The differences between wakefulness (frequently opened eyes, coordinated body movement and response to the environment) and deep sleep (closed eyes, lack of overt behaviour and absence of response to stimuli) are very well known. However, it has been shown that when wakefulness is prolonged, characteristic OFF periods of the slow wave of sleep can appear locally^87^. In the same way during sleep the frequency of these decreases as the sleep goes on. Therefore it would be expected that, at the level of the global brain in which we conducted our study, some of the characteristics of both states are shared while the differences have to be established in a statistical way. Indeed, if we consider the medium-high values of *q* characteristic of the sleeping condition, we see that the awake brain can reach these values during some intervals, analogously to a prolonged wakefulness with the characteristic OFF periods that can appear locally^87^.

The mean NoEL 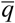, is higher when the participants are asleep as shown in Fig.7A. However, if we use as a reference the energy of the unstable trivial solution (0, 0,…, 0) and take into account that when the number of levels increases, the stability of the GASS grows and its energy decreases, the average energy of the deep asleep state is less than the awake one.

Given that energy levels can be interpreted not only in terms of energy but also of attraction or stability (see section), this would indicate that, on average, the ISs found in deep sleep are more stable. The lower stability of the awake state can also be inferred from the greater variability of the NoEL in that state according to our results. All this agrees with other studies in which authors claim that, using very different methodologies, that the brain exhibits less stability during wakefulness ^80^.

More generally, the conscious state has usually been associated with complexity in different ways. Different measures of the dynamic complexity of a network have been proposed. For instance in^88^, given a network its pair-wise correlation matrix reflects the degree of interdependencies among the nodes. When the nodes are disconnected or close to independence (equivalent to a small *g*), no complex collective dynamics emerge and the distribution of cross-correlation values are characterized by a narrow peak close to low values. When the collective dynamics are close to global synchrony (equivalent to a large *g*) and the distribution of cross-correlation values has a peak near highest value. It is not a complex state either because all nodes follow the same behavior. Complexity emerges when the collective dynamics are characterized by intermediate states, between independence and global synchrony and it is characterized by a broad distribution of correlation values or interdependencies among the node values, so the *functional complexity* is reflected in the variability of the associated distribution. This variability can be defined as normed entropy or as the difference between distribution and uniform distribution quantified as the integral of the absolute value of the difference, for instance. So, dynamic complexity is clearly associated with more uniform distributions. Although our distributions are not made from cross-correlation values but from NoEL, the underlying idea is the same. The pattern of statistical distribution of NoEL in wakefulness (Fig.6C) can be interpreted as a sign of complexity. We have seen that in deep sleep the probability of a concrete *q* increases with *q*. On the contrary, when the participants are awake the frequency remains almost constant with *q*∈ [1, 89].

A greater unpredictability of the wakeful brain dynamics can also be considered because there is maximum entropy and minimal information about the state subsequent to the current one. Unpredictability is one of the features of complexity and chaotic dynamics. On the contrary, in deep sleep it is very likely to find the system in medium-high NoEL informational structure, restricting the repertoire of possible dynamics in this state.

In our results the complexity of the wakefulness state is reflected not in more complex ISs but in the variability of different ISs in this state. Taking the standard deviation as a measure of NoEL distribution variability, Fig.7B demonstrates that the functional complexity of an awake individual is higher (except for one case) than in the same individual during deep sleep.

It should be emphasised that although a fitting was made to obtain the optimal *g*, the usefulness of our method is not restricted to a narrow range of values of *g*, neither to a particular filtering range, as shown by the results reported in this study.

Our work can be interpreted as a method to calculate the empirical “ghost” attractors of the resting state suggested in^65^. These multi-stable attractors correspond to distinct foci of high firing activity in particular brain areas. But at the edge of certain bifurcation in the used ansatz they do not exist as stable fixed points yet since they are either saddle points, or regimes with close to zero flow in the phase space. However these states could be easily stabilized when needed in a given task context or for a given function. The concept of DIS serve here as the abstract object to describe multistability^77^ or metastability^89, 90^ and can be a useful tool to analyze the functional connectivity dynamics^91^.

The main contribution of this work is a solid theoretical framework for describing the attractor landscape of a brain state, validated with empirical neuroimaging data. In future, further steps in this regard include finding out what lies below this explanation of the complex dynamical landscape. The theoretical framework proposed is potentially highly relevant for the development of biomarkers in translational applications, especially in the context of patients with neuropsychiatric disorders and with different levels of coma.

## Methods: neuroimaging data

### Empirical data and filtering

In order to compare different naturally occurring brain states we use data from 18 healthy participants. Specifically, BOLD signals from fMRI in resting state and deep asleep phase (N3). Empirical data comes from a set of fifty-five subjects (thirty-six females, mean ±SD age of 23.4 ±3.3 years) who fell asleep during a simultaneous EEG-fMRI recording previously described in^92^, where 18 participants who reached stage N3 sleep (deep sleep) were selected. The mean duration (±standard deviation) of contiguous N3 sleep epochs for these participants was 11.67 ±8.66 min. fMRI data was recorded at 3T (Siemens Trio, Erlangen, Germany) simultaneously with EEG data using an MR-compatible EEG cap (modified BrainCapMR, Easycap, Herrsching, Germany), MRI-compatible amplifiers (BrainAmp MR, BrainProducts, Garching, Germany), and sleep stages were scored manually by an expert according to the AASM criteria (AASM, 2007). fMRI data was realigned, normalized and spatially smoothed using SPM8 (www.fil.ion.ucl.ac.uk/spm), (see Supplementary Information and^92^ for full acquisition, pre-processing and sleep scoring details). Cardiac (with frequencies around 1∼2 Hz), respiratory (around 0.3 Hz), motion-induced and their temporal aliasing noises was regressed out from the fMRI BOLD signals^93^, and data was band-pass filtered, first in the range 0.04−0.07 Hz, and then in other ranges^94^.

Empirical BOLD signals are normally band-pass filtered to remove the contribution of noise. In this study we take a particular look at this process to better illustrate how instantaneous attractors found in resting state and in the deep sleep differentiate from one another. Different filters have been proposed in the literature to obtain the most reliable results. For instance, the range 0.04−0.07 Hz is ideal according to^95^. We consider all the possible filters below the Nyquist frequency to search of the one that best fits the purpose of showing how instantaneous attractors in awake and asleep conditions differentiate.

### Structural connectivity matrix

For the whole-brain network model, the interactions between the 90 brain areas were scaled in proportion to their white matter structural connectivity. For this study, we used the structural connectivity between the 90 automated anatomical labelling (AAL) regions obtained in a previous study^70^ averaged across 16 healthy young adults (5 females, mean ± SD age: 24.75±2.54). Briefly, for each subject, a 90×90 structural connectivity matrix Γ=[*γ*_*i j*_] was obtained by applying tractography algorithms to Diffusion magnetic resonance imaging (dMRI) following the same methodology described in^96^ where the connectivity *γ*_*i j*_ between regions *i* and *j* is calculated as the proportion of sampled fibers in all voxels in region *i* that reach any voxel in region *j*. Since dMRI does not capture fiber directionality, *γ*_*i j*_ was defined as the average between *γ*_*i j*_ and *γ* _*ji*_. Averaging across all 16 participants resulted in a structural connectivity matrix Γ=[*γ*_*i j*_] representative of healthy young adults. We order the different brain areas in the neuroanatomical connectivity matrix in such a way that homotopic regions in the two cerebral hemispheres were arranged symmetrically with respect to the center of the matrix(see Supplementary Information including Fig. 2.1 for more details).

## Acknowledgements

J.A.G., I.G. and J.A.L have been partially supported by Spanish Ministerio de Economãa y Competitividad and FEDER, projects MTM2015-63723-P, PGC2018-096540-B-I00, and Proyecto I+D+i Programa Operativo FEDER Andalucãa US-1254251. S.S.P. is supported by the Economic and Social Research Council (UK) under the project ES/S010947/1. G.D. is supported by the Spanish Research Project PSI2016-75688-P (Agencia Estatal de Investigacion/Fondo Europeo de DesarrolloRegional, European Union); by the European Union’s Horizon 2020 Research and Innovation Programme under Grant Agreements 720270 (Human Brain Project [HBP] SGA1) and 785907 (HBP SGA2); and by the Catalan Agency for Management of University and Research Grants Programme 2017 SGR 1545. M.L.K. is supported by the European Research Council Consolidator Grant: CAREGIVING (615539) and Center for Music in the Brain, funded by the Danish National Research Foundation (DNRF117). H.L. and data acquisition were supported by the Bundesministerium für Bildung und Forschung (grant no. 01 EV 0703); H.L. and E.T. by the LOEWE Neuronale Koordination Forschungsschwerpunkt Frankfurt (NeFF); H.L. also by Christian-Albrechts-University Kiel and the CRC1261 (Deutsche Forschungsgemeinschaft). We thank Astrid Morzelewski for sleep scoring of the EEGs.

## Author contributions statement

M.L.K. provided SCM measurements. H.L. designed and supervised the EEG-fMRI experiments including EEG and BOLD data (pre-)processing together with E.T. and sleep scoring. J.A.G., S.S.P., I.G., J.A.L. and G.D. did the theoretical modelling. J.A.G. and S.S.P. with help from M.L.K created all the figures. J.A.G performed calculations and wrote the majority of the text. G.D. and J.A.L. were the lead principal investigators. All authors contributed significantly to discussions, to the editing of the text and reviewed the final manuscript.

## References

1. Deco, G. et al. Awakening: predicting external stimulation forcing transitions between different brain states. PNAS 116, 18088–97 (2019).

2. Casali, A. G. et al. A theoretically based index of consciousness independent of sensory processing and behavior. Sci Transl Med 5, 198ra105, DOI: 10.1126/scitranslmed.3006294 (2013).

3. Casali, Adenauer G Gosseries, Olivia Rosanova, Mario Boly, Melanie Sarasso, Simone Casali, Karina R Casarotto, Silvia Bruno, Marie-Aurelie Laureys, Steven Tononi, Giulio Massimini, Marcello eng Research Support, Non-U.S. Gov’t 2013/08/16 06:00 Sci Transl Med. 2013 Aug 14;5(198):198ra105. DOI: 10.1126/scitranslmed.3006294.

3. Oizumi, M., Albantakis, L. & Tononi, G. From the phenomenology to the mechanisms of consciousness: Integrated information theory 3.0. PLOS Comput. Biol. 10, 1–25, DOI: 10.1371/journal.pcbi.1003588 (2014).

4. Tononi, G. An information integration theory of consciousness. BMC Neurosci. 5, 42, DOI: 10.1186/1471-2202-5-42 (2004).

5. Tononi, G. Consciousness as integrated information: a provisional manifesto. The Biol. Bull. 215, 216–242, DOI: 10.2307/25470707 (2008). PMID: 19098144, https://doi.org/10.2307/25470707.

6. Tononi, G. Integrated information theory of consciousness: an updated account. Arch. italiennes de biologie 150, 56—90, DOI: 10.4449/aib.v149i5.1388 (2012).

7. Tononi, G., Boly, M., Massimini, M. & Koch, C. Integrated information theory: from consciousness to its physical substrate. Nat. Rev. Neurosci. 17, 450 EP – (2016).

8. Pearl, J. Causality (Cambridge University Press, 2009).

9. Albantakis, L., Marshall, W., Hoel, E. P. & Tononi, G. What caused what? an irreducible account of actual causation. CoRR abs/1708.06716 (2017). 1708.06716.

10. Stevner, A. et al. Discovery of key whole-brain transitions and dynamics during human wakefulness and non-rem sleep. Nat. Commun. 10, 1035 (2019).

11. Ferrarelli, F. et al. Breakdown in cortical effective connectivity during midazolam-induced loss of consciousness. Proc. Natl. Acad. Sci. 107, 2681–2686, DOI: 10.1073/pnas.0913008107 (2010). https://www.pnas.org/content/107/6/2681.full.pdf.

12. Rosanova, M. et al. Recovery of cortical effective connectivity and recovery of consciousness in vegetative patients. Brain 135, DOI: 10.1093/brain/awr340 (2012). http://oup.prod.sis.lan/brain/article-pdf/135/4/1308/17869827/awr340.pdf.

13. Carhart-Harris, R. et al. The entropic brain: a theory of conscious states informed by neuroimaging research with psychedelic drugs. Front. Hum. Neurosci. 8, 20, DOI: 10.3389/fnhum.2014.00020 (2014).

14. Esteban, F. J., Galadí, J. A., Langa, J. A., Portillo, J. R. & Soler-Toscano, F. Informational structures: A dynamical system approach for integrated information. PLOS Comput. Biol. 14, 1–33, DOI: 10.1371/journal.pcbi.1006154 (2018).

15. Kalita, P., Langa, J. A. & Soler-Toscano, F. Informational structures and informational fields as a prototype for the description of postulates of the integrated information theory. Entropy 21, DOI: 10.3390/e21050493 (2019).

16. Bassett, D. S. & Bullmore, E. Small-world brain networks. The Neurosci. 12, 512–523, DOI: 10.1177/1073858406293182 (2006). PMID: 17079517, https://doi.org/10.1177/1073858406293182.

17. Markov, N. T. et al. Cortical high-density counterstream architectures. Science 342, 1238406, DOI: 10.1126/science.1238406 (2013).

19. Markov, Nikola T Ercsey-Ravasz, Maria Van Essen, David C Knoblauch, Kenneth Toroczkai, Zoltan Kennedy, Henry eng R01 MH060974/MH/NIMH NIH HHS/ R01 MH60974/MH/NIMH NIH HHS/ Research Support, N.I.H., Extramural Research Support, Non-U.S. Gov’t Research Support, U.S. Gov’t, Non-P.H.S. Review New York, N.Y. 2013/11/02 06:00 Science. 2013 Nov 1;342(6158):1238406. doi: 10.1126/science.1238406.

18. Guerrero, G., Langa, J. A. & Suárez, A. Attracting complex networks. In Complex networks and dynamics, vol. 683 of Lecture Notes in Econom. and Math. Systems, 309–327 (Springer, [Cham], 2016).

19. Guerrero, G., Langa, J. A. & Suárez, A. Architecture of attractor determines dynamics on mutualistic complex networks. Nonlinear Anal. Real World Appl. 34, 17–40, DOI: 10.1016/j.nonrwa.2016.07.009 (2017).

20. Bascompte, J. & Jordano, P. Mutualistic Networks. Monographs in population biology (Princeton University Press, 2014).

21. Bastolla, U. et al. The architecture of mutualistic networks minimizes competition and increases biodiversity. Nature 458, 1018–1020 (2009).

22. Bascompte, J., Jordano, P., Melián, C. J. & Olesen, J. M. The nested assembly of plant–animal mutualistic networks. Proc. Natl. Acad. Sci. 100, 9383–9387, DOI: 10.1073/pnas.1633576100 (2003). http://www.pnas.org/content/100/16/9383.full.pdf.

23. Bascompte, J. & Jordano, P. Plant-animal mutualistic networks: The architecture of biodiversity. Annu. Rev. Ecol. Evol. Syst. 38, 567–593, DOI: 10.1146/annurev.ecolsys.38.091206.095818 (2007). https://doi.org/10.1146/annurev.ecolsys.38.091206.095818.

24. Barrat, A., Barthélemy, M. & Vespignani, A. Dynamical Processes on Complex Networks (Cambridge University Press, 2008).

25. Gilarranz, L. J., Rayfield, B., Liñán-Cembrano, G., Bascompte, J. & Gonzalez, A. Effects of network modularity on the spread of perturbation impact in experimental metapopulations. Science 357, 199–201, DOI: 10.1126/science.aal4122 (2017). https://science.sciencemag.org/content/357/6347/199.full.pdf.

26. Rohr, R. P., Saavedra, S. & Bascompte, J. On the structural stability of mutualistic systems. Science 345, DOI: 10.1126/science.1253497 (2014). https://science.sciencemag.org/content/345/6195/1253497.full.pdf.

27. Golos, M., Jirsa, V. & Dauc′, E. Multistability in large scale models of brain activity. PLOS Comput. Biol. 11, 1–32, DOI: 10.1371/journal.pcbi.1004644 (2016).

28. Deco, G., Senden, M. & Jirsa, V. How anatomy shapes dynamics: a semi-analytical study of the brain at rest by a simple spin model. Front. Comput. Neurosci. 6, 68, DOI: 10.3389/fncom.2012.00068 (2012).

29. Puu, T. Attractors, Bifurcations, and Chaos: Nonlinear Phenomena in Economics (Springer Berlin Heidelberg, 2013).

30. Nir, Y., Massimini, M., Boly, M. & Tononi, G. Sleep and Consciousness, 133–182 (Springer Berlin Heidelberg, Berlin, Heidelberg, 2013).

31. Stevner, A. et al. Discovery of key whole-brain transitions and dynamics during human wakefulness and non-rem sleep. Nat. Commun. 10, 1035 (2019).

32. Strogatz, S. Nonlinear Dynamics And Chaos. Studies in nonlinearity (Sarat Book House, 2007).

33. Wiggins, S. Introduction to Applied Nonlinear Dynamiccal Systems and Chaos. Introduction to Applied Nonlinear Dynamiccal Systems and Chaos, (Springer-Verlag New York, I., 2003).

34. Glendinning, P. Stability, Instability and Chaos: An Introduction to the Theory of Nonlinear Differential Equations. Cambridge Texts in Applied Mathematics (Cambridge University Press, 1994).

35. Arnol’d, V. Catastrophe Theory (Springer Berlin Heidelberg, 2003).

36. Hirsch, M., Smale, S. & Devaney, R. Differential Equations, Dynamical Systems, and an Introduction to Chaos (Elsevier Science, 2012).

37. Sandefur, J. Discrete Dynamical Systems: Theory and Applications (Clarendon Press, 1990).

38. Babin, A. & Vishik, M. Attractors of Evolution Equations. Studies in Mathematics and its Applications (Elsevier Science, 1992).

39. Hale, J. Asymptotic Behavior of Dissipative Systems. Mathematical surveys and monographs (American Mathematical Society, 1988).

40. Henry, D. B. Geometric theory of semilinear parabolic equations (Springer-Verlag, Berlin, 1981).

41. Ladyzhenskaya, O. A. Attractors for semigroups and evolution equations (Cambridge University Press, 1991).

42. Robinson, J., Crighton, D. & Ablowitz, M. Infinite-Dimensional Dynamical Systems: An Introduction to Dissipative Parabolic PDEs and the Theory of Global Attractors. Cambridge Texts in Applied Mathematics (Cambridge University Press, 2001).

43. Temam, R. Infinite-Dimensional Dynamical Systems in Mechanics and Physics. Applied Mathematical Sciences (Springer New York, 1997).

44. Aragao-Costa, E. R., Caraballo, T., Carvalho, A. N. & Langa, J. A. Stability of gradient semigroups under perturbations. Nonlinearity 24, 2099 (2011).

45. Conley, C. Isolated invariant sets and the Morse index. No. 38 in CBMS Regional Conference Series in Mathematics (American Mathematical Society, Providence, 1978).

46. Babin, A. V. & Vishik, M. Regular attractors of semigroups and evolution equations. Math. Pures et Appl. 62, 441–491 (1983).

47. Carvalho, A., Langa, J. & Robinson, J. Attractors for infinite-dimensional non-autonomous dynamical systems. Applied Mathematical Sciences (Springer New York, 2012).

48. Norton, D. E. The fundamental theorem of dynamical systems. Commentationes Math. Univ. Carol. 36, 585–597 (1995).

49. Aragao-Costa, E. R., Caraballo, T., Carvalho, A. N. & Langa, J. A. Continuity of lyapunov functions and of energy level for a generalized gradient semigroup. Topol. Methods Nonlinear Anal. 39, 57–82 (2012).

50. Joly, R. & Raugel, G. Genertic morse-smale property for the parabolic equation on the circle. Annales de l’Institut Henri Poincare (C) Non Linear Analysis 27, 1397–1440, DOI: https://doi.org/10.1016/j.anihpc.2010.09.001 (2010).

51. Hale, J., Magalhaes, L. & Oliva, W. An Introduction to Infinite Dimensional Dynamical Systems -Geometric Theory. Applied Mathematical Sciences (Springer New York, 2013).

52. Palis, J., Manning, A. & de Melo, W. Geometric Theory of Dynamical Systems: An Introduction (Springer New York, 2012).

53. Ott, E. Chaos in Dynamical Systems (Cambridge University Press, 2002).

54. Porter, M. & Gleeson, J. Dynamical Systems on Networks: A Tutorial. Frontiers in Applied Dynamical Systems: Reviews and Tutorials (Springer International Publishing, 2016).

55. Mortveit, H. & Reidys, C. An Introduction to Sequential Dynamical Systems (Springer US, 2008).

56. Osipenko, G. Dynamical Systems, Graphs, and Algorithms. Lecture Notes in Mathematics (Springer Berlin Heidelberg, 2006).

57. Murray, J. Mathematical Biology. Biomathematics (Springer Berlin Heidelberg, 2013).

58. Takeuchi, Y. Global Dynamical Properties of Lotka-Volterra Systems (World Scientific, 1996).

59. Patrão, M. & San Martin, L. A. B. Semiflows on topological spaces: chain transitivity and semigroups. J. Dynam. Differ. Equations 19, 155–180, DOI: 10.1007/s10884-006-9032-3 (2007).

60. Afraimovich, V. S., Moses, G. & Young, T. Two-dimensional heteroclinic attractor in the generalized Lotka-Volterra system. Nonlinearity 29, 1645, DOI: 10.1088/0951-7715/29/5/1645 (2016). 1509.04570.

61. Afraimovich, V. S., Zhigulin, V. P. & Rabinovich, M. I. On the origin of reproducible sequential activity in neural circuits. Chaos: An Interdiscip. J. Nonlinear Sci. 14, 1123–1129, DOI: 10.1063/1.1819625 (2004). https://doi.org/10.1063/1.1819625.

62. Afraimovich, V., Tristan, I., Varona, P. & Rabinovich, M. Transient dynamics in complex systems: Heteroclinic sequences with multidimensional unstable manifolds. Discontinuity, Nonlinearity, Complex. 2(1), 21–41, DOI: 10.5890/DNC.2012.11.001 (2013).

63. Muezzinoglu, M. K., Tristan, I., Huerta, R., Afraimovich, V. S. & Rabinovich, M. I. Transients versus attractors in complex networks. Int. J. Bifurc. Chaos 20, 1653–1675, DOI: 10.1142/S0218127410026745 (2010). https://doi.org/10.1142/S0218127410026745.

64. Rabinovich, M., Varona, P., Tristan, I. & Afraimovich, V. Chunking dynamics: heteroclinics in mind. Front. Comput. Neurosci. 8, 22, DOI: 10.3389/fncom.2014.00022 (2014).

65. Deco, G. & Jirsa, V. K. Ongoing cortical activity at rest: Criticality, multistability, and ghost attractors. J. Neurosci. 32, 3366–3375, DOI: 10.1523/JNEUROSCI.2523-11.2012 (2012). http://www.jneurosci.org/content/32/10/3366.full.pdf.

66. Cottle, R., Pang, J. & Stone, R. The Linear Complementarity Problem. Classics in Applied Mathematics (Society for Industrial and Applied Mathematics (SIAM, 3600 Market Street, Floor 6, Philadelphia, PA 19104), 1992).

67. Murty, K. Linear Complementarity, Linear and Non Linear Programming. Sigma series in applied mathematics (Heldermann Verlag, 1988).

68. Cross, G. Three types of matrix stability. Linear Algebr. its Appl. 20, 253 – 263, DOI: https://doi.org/10.1016/0024-3795(78)90021-6 (1978).

69. Takeuchi, Y. & Adachi, N. The existence of globally stable equilibria of ecosystems of the generalized volterra type. J. Math. Biol. 10, 401–415, DOI: 10.1007/BF00276098 (1980).

70. Deco, G. et al. Single or multiple frequency generators in on-going brain activity: A mechanistic whole-brain model of empirical meg data. NeuroImage 152, 538 – 550, DOI: https://doi.org/10.1016/j.neuroimage.2017.03.023 (2017).

71. Cammoun, L. et al. Mapping the human connectome at multiple scales with diffusion spectrum mri. J. Neurosci. Methods 203, 386 – 397, DOI: https://doi.org/10.1016/j.jneumeth.2011.09.031 (2012).

72. Collins, D. L. et al. Design and construction of a realistic digital brain phantom. IEEE Transactions on Med. Imaging 17, 463–468, DOI: 10.1109/42.712135 (1998).

73. Tzourio-Mazoyer, N. et al. Automated anatomical labeling of activations in spm using a macroscopic anatomical parcellation of the mni mri single-subject brain. NeuroImage 15, 273 – 289, DOI: https://doi.org/10.1006/nimg.2001.0978 (2002).

74. Cabral, J., Kringelbach, M. L. & Deco, G. Exploring the network dynamics underlying brain activity during rest. Prog. Neurobiol. 114, 102 – 131, DOI: https://doi.org/10.1016/j.pneurobio.2013.12.005 (2014).

75. Deco, G., Jirsa, V., McIntosh, A. R., Sporns, O. & Kötter, R. Key role of coupling, delay, and noise in resting brain fluctuations. Proc. Natl. Acad. Sci. 106, 10302–10307, DOI: 10.1073/pnas.0901831106 (2009). https://www.pnas.org/content/106/25/10302.full.pdf.

76. Deco, G., Tononi, G., Boly, M. & Kringelbach, M. L. Rethinking segregation and integration: contributions of whole-brain modelling. Nat. Rev. Neurosci. 16, 430 EP – (2015).

77. Kelso, J. A. S. Multistability and metastability: understanding dynamic coordination in the brain. Philos. Transactions Royal Soc. B: Biol. Sci. 367, 906–918, DOI: 10.1098/rstb.2011.0351 (2012).

78. Yorke, J. A. & Anderson, W. N. Predator-prey patterns. Proc. Natl. Acad. Sci. United States Am. 70, 2069–2071, DOI: 10.1073/pnas.70.7.2069 (1973).

79. Mueller, S. et al. Individual variability in functional connectivity architecture of the human brain. Neuron 77, 586 – 595, DOI: https://doi.org/10.1016/j.neuron.2012.12.028 (2013).

80. Jobst, B. M. et al. Increased stability and breakdown of brain effective connectivity during slow-wave sleep: Mechanistic insights from whole-brain computational modelling. Sci. Reports 7, 4634, DOI: 10.1038/s41598-017-04522-x (2017).

81. Boly, M. et al. Hierarchical clustering of brain activity during human nonrapid eye movement sleep. Proc. Natl. Acad. Sci. 109, 5856–5861, DOI: 10.1073/pnas.1111133109 (2012). https://www.pnas.org/content/109/15/5856.full.pdf.

82. Tagliazucchi, E. et al. Breakdown of long-range temporal dependence in default mode and attention networks during deep sleep. Proc. Natl. Acad. Sci. 110, 15419–15424, DOI: 10.1073/pnas.1312848110 (2013). https://www.pnas.org/content/110/38/15419.full.pdf.

83. Spoormaker, V. I. et al. Development of a large-scale functional brain network during human non-rapid eye movement sleep. J. Neurosci. 30, 11379–11387, DOI: 10.1523/JNEUROSCI.2015-10.2010 (2010). http://www.jneurosci.org/content/30/34/11379.full.pdf.

84. Spoormaker, V., Gleiser, P. & Czisch, M. Frontoparietal connectivity and hierarchical structure of the brain’s functional network during sleep. Front. Neurol. 3, 80, DOI: 10.3389/fneur.2012.00080 (2012).

85. Dehaene, S., Charles, L., King, J.-R. & Marti, S. Toward a computational theory of conscious processing. Curr. Opin. Neurobiol. 25, 76 – 84, DOI: https://doi.org/10.1016/j.conb.2013.12.005 (2014). Theoretical and computational neuroscience.

86. Alkire, M. T., Hudetz, A. G. & Tononi, G. Consciousness and anesthesia. Science 322, 876–880, DOI: 10.1126/science.1149213 (2008). https://science.sciencemag.org/content/322/5903/876.full.pdf.

87. Vyazovskiy, V. V. et al. Local sleep in awake rats. Nature 472, >443 EP – (2011).

88. Zamora-López, G., Chen, Y., Deco, G., Kringelbach, M. L. & Zhou, C. Functional complexity emerging from anatomical constraints in the brain: the significance of network modularity and rich-clubs. Sci. Reports 6, 38424 EP – (2016).

89. Tognoli, E. & Kelso, J. The metastable brain. Neuron 81, 35 – 48, DOI: https://doi.org/10.1016/j.neuron.2013.12.022 (2014).

90. Werner, G. Metastability, criticality and phase transitions in brain and its models. Biosystems 90, 496 – 508, DOI: https://doi.org/10.1016/j.biosystems.2006.12.001 (2007).

91. Hansen, E. C., Battaglia, D., Spiegler, A., Deco, G. & Jirsa, V. K. Functional connectivity dynamics: Modeling the switching behavior of the resting state. NeuroImage 105, 525 – 535, DOI: https://doi.org/10.1016/j.neuroimage.2014.11.001 (2015).

92. Tagliazucchi, E. & Laufs, H. Decoding wakefulness levels from typical fmri resting-state data reveals reliable drifts between wakefulness and sleep. Neuron 82, 695 – 708, DOI: https://doi.org/10.1016/j.neuron.2014.03.020 (2014).

93. Glover, G. H., Li, T.-Q. & Ress, D. Image-based method for retrospective correction of physiological motion effects in fmri: Retroicor. Magn. Reson. Medicine 44, 162–167, DOI: 10.1002/1522-2594(200007)44:1<162::AID-MRM23>3.0.CO;2-E (2000).

94. Cordes, D. et al. Frequencies contributing to functional connectivity in the cerebral cortex in “resting-state” data. Am. J. Neuroradiol. 22, 1326–1333 (2001). http://www.ajnr.org/content/22/7/1326.full.pdf.

95. Glerean, E., Salmi, J., Lahnakoski, J. M., Jääskeläinen, I. P. & Sams, M. Functional magnetic resonance imaging phase synchronization as a measure of dynamic functional connectivity. Brain Connect. 2, 91–101, DOI: 10.1089/brain.2011.0068 (2012). PMID: 22559794, https://doi.org/10.1089/brain.2011.0068.

96. Cabral, J., Kringelbach, M. L. & Deco, G. Functional graph alterations in schizophrenia: A result from a global anatomic decoupling? Pharmacopsychiatry 45, S57–S64, DOI: 10.1055/s-0032-1309001 (2012).

