## Supplementary Information for "Dynamical informational structures characterize the different human brain states of wakefulness and deep sleep"


#### ABSTRACT

#### 1 Supplementary information about figures

##### 1.1 Figure 1

$\alpha = (0.71, 2.85)$

$$g\Gamma = \begin{pmatrix} 0 & 0.1 \\ 0.2 & 0 \end{pmatrix}$$

##### 1.2 Figure 2

$$g\Gamma = \begin{pmatrix} 0 & 0.238218 & 0.111796 & 0.195597 \\ 0.163211 & 0 & 0.0450021 & 0.223199 \\ 0.141685 & 0.0952128 & 0 & 0.182694 \\ 0.109297 & 0.18679 & 0.0118409 & 0 \end{pmatrix}$$

At $t = 22$ there are 2 energy levels, at $t = 76$ there are 3 energy levels, at $t = 105$, $t = 122$, and $t = 150$ there are 4 levels of energy, and at $t = 54$ there are 5 energy levels. Always they match the number of non-zero variables of the globally asymptotically stable solution (GASS) plus one. The first level of energy (associated with a node or vertex with only outgoing edges) is always comprised of the trivial solution (all variables equal zero), while the last energy level includes only the GASS (associated with a node or vertex with only incoming edges). Depending on $\alpha$ and $g\Gamma$ the informational structure can be complete, i.e., including $2^n$ stationary points and the number of energy levels is $n+1$ (for instance, Fig.2.2).

#### 2 Supplementary models and methods

##### 2.1 Structural connectivity matrix

For the whole-brain network model, the interactions between the 90 brain areas were scaled in proportion to their white matter structural connectivity. For this study, we used the structural connectivity between the 90 AAL regions obtained in a previous study[1] averaged across 16 healthy young adults (5 females, mean $\pm$ SD age: 24.75 $\pm$ 2.54). Briefly, for each subject, a 90$\times$90 structural connectivity matrix $\Gamma = [\gamma_{ij}]$ was obtained by applying tractography algorithms to Diffusion Tensor Imaging (DTI) following the same methodology described in[2] where the connectivity $\gamma_{ij}$ between regions $i$ and $j$ is calculated as the proportion of sampled fibers in all voxels in region $i$ that reach any voxel in region $j$. Since DTI does not capture fiber directionality, $\gamma_{ij}$ was defined as the average between $\gamma_{ij}$ and $\gamma_{ji}$. Averaging across all 16 participants resulted in a structural connectivity matrix $\Gamma = [\gamma_{ij}]$ representative of healthy young adults. We order the different brain areas in the neuroanatomical connectivity matrix in such a way that homotopic regions in the two cerebral hemispheres were arranged symmetrically with respect to the center of the matrix.

##### 2.2 The Informational Structure (IS) of a dynamical system

###### *2.2.1 Global Attractor*

The phase space $X$ (for example $X$ could be $\mathbb{R}^N$) represents the framework in which the dynamics described by a semigroup of transformations $S(t) : X \to X$ is developed. Given a *phase space* $X$ we define a dynamical system (DS) on $X$ as a flow

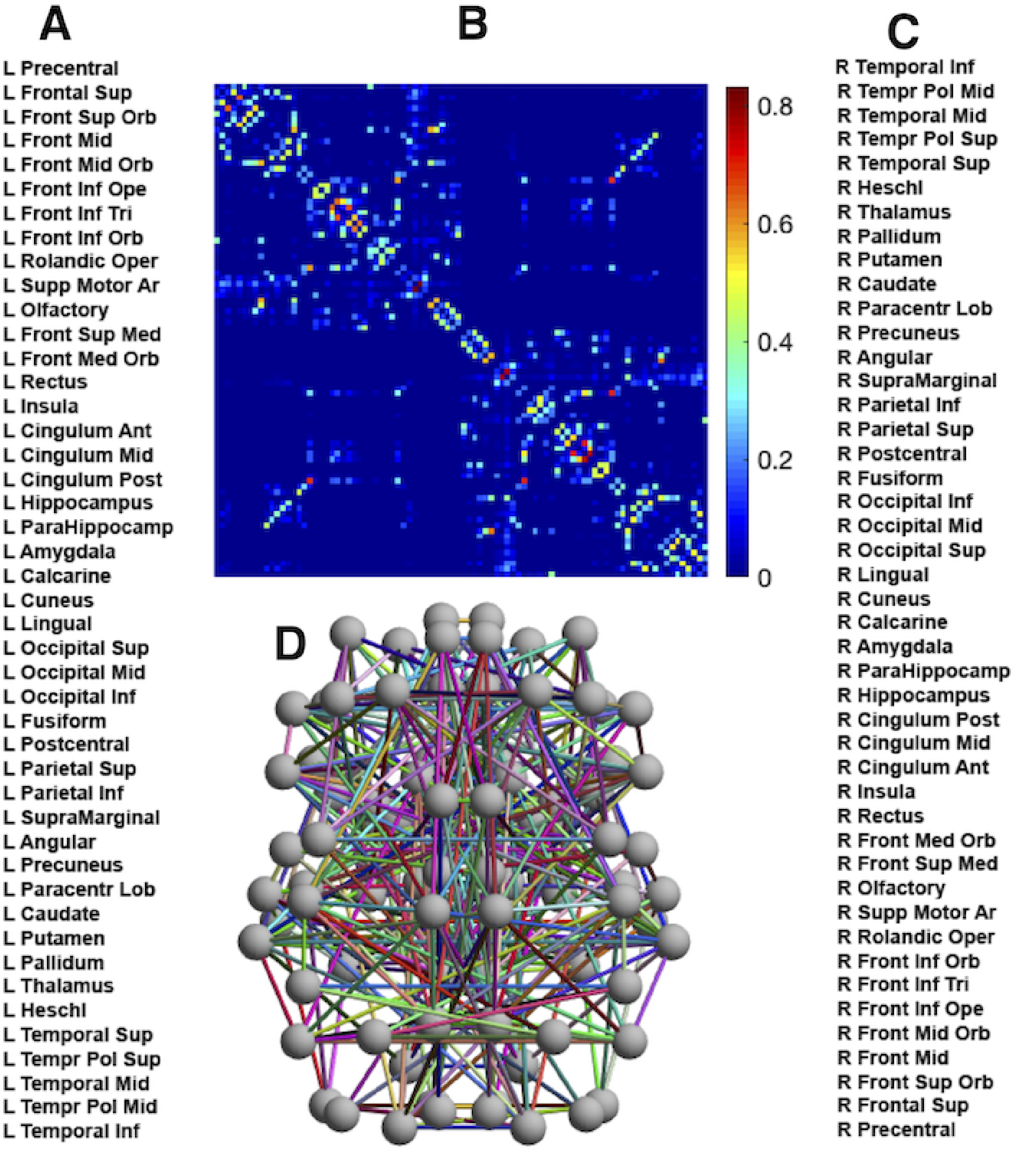


**Figure 2.1.** Structural connectivity of the Human Connectome. **A**, Cortical regions index of 90 regions from the Automated Anatomical Labeling (AAL) atlas with region labels on the left side and (**C**) homotopic regions on the right side. **B**, Structural connectivity matrix. Contra-lateral (homotopic) regions are symmetrically arranged. The anti-diagonal reveals the existing connections between contra-lateral regions. **D**, 2-dimensional representation of the network structure (view from above), the nodes representing anatomical regions placed at their central coordinates.

or family of non-linear operators $\{S(t)\}_{t\in\mathbb{R}^+}$, where $S(t)u \in X$ describes the dynamics of each element $u \in X$. In particular, $u(t;u_0) = S(t)u_0$ is a solution of the DS at time $t$ with initial condition $u_0$. Although $t$ is positive ($t \in \mathbb{R}^+$) there are solutions for $s \in \mathbb{R}$: a general or global solution associated to $S(t)$ would be $\xi : \mathbb{R} \to X$ such that $\xi(t+s) = S(t)\xi(s)$ for all $s \in \mathbb{R}$, $t \in \mathbb{R}^+$.

The Global Attractor ($\mathscr{GA}$) describes all the past and future scenarios of a DS and is an important concept in Dynamical Systems Theory. The $\mathscr{GA}$ is defined as follows[3–8]:

**Definition.** A set $\mathscr{A} \subseteq X$ is a $\mathscr{GA}$ for $\{S(t) : t \geq 0\}$ if it is

(i) compact,

(ii) invariant under $\{S(t) : t \geq 0\}$, i.e. $S(t)\mathscr{A} = \mathscr{A}$ for all $t \geq 0$, and

(iii) attracts bounded subsets of $X$ under $\{S(t) : t \geq 0\}$ for the Hausdorff semidistance; i.e., for all bounded $B \subset X$

$$\lim_{t\to+\infty} \mathrm{dist}_H(S(t)B, \mathscr{A}) := \lim_{t\to+\infty} \sup_{b\in B} \inf_{a\in\mathscr{A}} d(S(t)b, a) = 0.$$

Observe that (ii) can be interpreted as showing that an attractor has a proper intrinsic dynamics. Moreover, (iii) points that this set is determining all the future dynamics on the phase space $X$.

**Definition.** We say that $u^* \in X$ is an equilibrium point (or stationary solution) for the semigroup $S(t)$ if $S(t)u^* = u^*$, for all $t \geq 0$.

A stationary point is the simplest instance of global solution associated with $S(t)$. Futhermore stationary points are the minimal invariant objects inside a $\mathscr{GA}$. It is easy to show that every invariant set is a subset of the $\mathscr{GA}$[4].

Moreover, the $\mathscr{GA}$ has the following properties[7,8]:

(a) It is the maximal invariant set in the phase space.

(b) It is the smallest closed attracting set.

(c) It is made of bounded complete solutions, i.e., solutions that exists for all time $t \in \mathbb{R}$, and so giving information for the asymptotic past of the system.

###### 2.2.2 Informational Structure: a definition

Generically, the $\mathscr{GA}$ structure can be described by isolated invariant sets (typically stationary points, periodic orbits[9–12], but also chaotic dynamics[13–15]) and connecting global solutions among them[9,16]. Those connections among invariant sets describe its structure[17,18].

Let us see some preliminary concepts:

**Definition.** An undirected graph is an ordered pair $\mathscr{G} = (V, L)$ comprising a non-empty set $V$ of vertices (or nodes) together with a set $L$ of unordered pairs or edges joining 2-element of $V$. A directed graph or digraph is a graph in which edges (named arcs) have orientations, that is to say, edges are ordered pairs: $L \subset V \times V$.

In all the following definitions we will consider that a DS on $X$ with semigroup $\{S(t) : t \geq 0\}$ is given.
A global solution $\xi_{A,B} : \mathbb{R} \to X$ connects a set $A$ to other set $B$ if and only if
$\lim_{t\to-\infty} dist(\xi_{A,B}(t), A) = 0$ and $\lim_{t\to+\infty} dist(\xi_{A,B}(t), B) = 0$.

**Definition.** Two sets are connected when there is a global solution that fulfills the above. If there is no such global solution, they are not connected. $\mathscr{O}_\varepsilon(\Xi)$ is called uniform neighbourhood with radius $\varepsilon > 0$ of a set $\Xi$ if for all elements $u \in \Xi$, $B_\varepsilon(u) := \{x \in X \mid d(x,u) < \varepsilon\}$ is contained in $\mathscr{O}_\varepsilon(\Xi)$.

We say that an invariant set $\Xi \subset Z$ is an isolated invariant set if there is an $\varepsilon > 0$ such that $\Xi$ is the maximal invariant subset of $\mathscr{O}_\varepsilon(\Xi)$.

**Definition.** A disjoint family of isolated invariant sets is a family $\{\Xi_1, ..., \Xi_N\}$ of isolated invariant sets with the property that, $\mathscr{O}_\varepsilon(\Xi_i) \bigcap \mathscr{O}_\varepsilon(\Xi_j) = \varnothing$ , $1 \leqslant i < j \leqslant N$ for some $\varepsilon > 0$.

**Definition.** Consider a DS with a disjoint family of isolated invariant sets $\Omega = \{\Xi_1, ..., \Xi_N\}$. Let

$$\delta_0 = \frac{1}{2} \min_{1\leqslant i<j\leqslant N} d(\Xi_i, \Xi_j) > 0$$

Let $\varepsilon_0 < \delta_0$, $\Xi \in \Omega$ and $\varepsilon \in (0, \varepsilon_0)$. An $\varepsilon$-chain from $\Xi$ to $\Xi$ is a subset $\{\Xi_{l_1}, ..., \Xi_{l_k}\}$ of $\Omega$, together with points $\{y_1, ..., y_k\}$ in $X$ and $\{t_1, \sigma_1, ..., t_k, \sigma_k\}$ in $\mathbb{R}$ such that, $0 < \sigma_i < t_i$, $1 \leqslant i \leqslant k$, $k \leqslant n$, $d(y_i, \Xi_{l_i}) < \varepsilon$, $1 \leqslant i \leqslant k$, $\Xi = \Xi_{l_1} = \Xi_{l_{k+1}}$, $d(T(\sigma_i)y_i, \bigcup_{i=1}^{n} \Xi_i) > \varepsilon_0$ and $d(T(t_i)y_i, \Xi_{l_{i+1}}) < \varepsilon$, $1 \leqslant i \leqslant k$.

**Definition.** We say that $\Xi \in \Xi$ is chain recurrent if there exist a fixed $\varepsilon_0 > 0$ and an $\varepsilon$-chain from $\Xi$ to $\Xi$, for each $\varepsilon \in (0, \varepsilon_0)$.

**Definition.** Let $\Omega = \{\Xi_1, ..., \Xi_N\}$ a disjoint family of isolated invariant sets and assume that it possesses a $\mathscr{G}\mathscr{A}$ $\mathscr{A}$. We say that $\{S(t) : t \geqslant 0\}$ is a generalized gradient-like semigroup relative to $\Xi$ if the following conditions are satisfied:

(G1) For any global solution $\xi : \mathbb{R} \to \mathscr{A}$, there are $1 \leqslant i, j \leqslant N$ such that

$$\lim_{t\to-\infty} dist(\xi(t), \Xi_i) = 0$$

and

$$\lim_{t\to\infty} dist(\xi(t), \Xi_j) = 0.$$

(G2) $\Omega = \{\Xi_1, ..., \Xi_N\}$ has no chain recurrent sets.

**Definition.** When each $\Xi_i$ consists only of a single stationary point, we say that the semigroup is a gradient-like semigroup.

**Definition.** A homoclinic structure associated with $\Omega = \{\Xi_1, ..., \Xi_N\}$ is a subset $\{\Xi_{l_1}, ..., \Xi_{l_k}\}$ of $\Omega$ ($k \leqslant N$) together with a set of global solutions $\{\xi_j,\ 1 \leq j \leq k\}$ such that

$$\lim_{t\to-\infty} dist(\xi_j(t), \Xi_{l_j}) = 0$$

and

$$\lim_{t\to\infty} dist(\xi_j(t), \Xi_{l_{j+1}}) = 0$$

for $1 \leq j \leq k$, and $\Xi_{l_{k+1}} := \Xi_{l_1}$.

**Theorem.** *Let $\{S(t) : t \geqslant 0\}$ be a semigroup with a disjoint family of isolated invariant sets $\Omega = \{\Xi_1, ..., \Xi_N\}$ and a global attractor $\mathscr{A}$. If $\{S(t) : t \geqslant 0\}$ satisfies (G1), then (G2) is satisfied if and only if $\mathscr{A}$ has no homoclinic structures.*

With this result we can redefine the concept of generalized gradient-like semigroups in the following equivalent way[16]:

**Definition.** Let $\{S(t) : t \geqslant 0\}$ be a semigroup with a disjoint family of isolated invariant sets $\Omega = \{\Xi_1, ..., \Xi_N\}$ and a $\mathscr{G}\mathscr{A}$ $\mathscr{A}$. We say that $\{S(t) : t \geqslant 0\}$ is a generalized gradient-like semigroup relative to $\Omega$ if:

(G1) For any global solution $\xi : \mathbb{R} \to \mathscr{A}$, there are $1 \leqslant i, j \leqslant N$ such that

$$\lim_{t\to-\infty} dist(\xi(t), \Xi_i) = 0$$

and

$$\lim_{t\to\infty} dist(\xi(t), \Xi_j) = 0.$$

(G2´) There is no homoclinic structure associated to $\Omega$.

Next we introduce the notion of a Morse decomposition for the attractor $\mathscr{A}$ of a semigroup $\{S(t) : t \geqslant 0\}$. We start with the notion of attractor–repeller pair.

**Definition.** Let $\{S(t) : t \geqslant 0\}$ be a semigroup with a $\mathscr{GA}$ $\mathscr{A}$. We say that a non-empty subset $A$ of $\mathscr{A}$ is a local attractor if there is an $\varepsilon > 0$ such that $\omega(\mathscr{O}\varepsilon(A)) = A$. The repeller $A^*$ associated to a local attractor A is the set defined by

$$A^* := \left\{ x \in A : \omega(x) \bigcap A = \varnothing \right\}.$$

The pair $(A, A^*)$ is called attractor–repeller pair for $\{S(t) : t \geqslant 0\}$ . Note that if $A$ is a local attractor, then $A^*$ is closed and invariant.

**Definition.** Given an increasing family $\varnothing = A_0 \subset A_1 \subset \cdots \subset A_N = \mathscr{A}$, of $N+1$ local attractors, for $j = 1, ..., N$, define $\Xi_j := A_j \bigcap A^*_{j-1}$. The ordered $N$-tuple $\Omega = \{\Xi_1, ..., \Xi_N\}$ is called a Morse decomposition for $\mathscr{A}$[9,16].

An equivalent definition of Morse decomposition for the attractor $\mathscr{A}$ of a semigroup $\{S(t) : t \geqslant 0\}$ is the following:

**Definition.** Let $\{S(t) : t \geqslant 0\}$ be a semigroup with a $\mathscr{GA}$ $\mathscr{A}$. Assume that there exists a collection $\Omega = \{\Xi_1, ..., \Xi_N\}$ of disjoint, compact and invariant subsets of $\mathscr{A}$ satisfying the following: for a given global solution $\xi : \mathbb{R} \to \mathscr{A}$

1. either $\xi(t) \in \Xi_i$, for all $t \in \mathbb{R}$ and some $i = 1, ..., N$;
2. or there exist $1 \leqslant i, j \leqslant N$ such that

$$\lim_{t \to -\infty} dist(\xi(t), \Xi_i) = 0$$

and

$$\lim_{t \to \infty} dist(\xi(t), \Xi_j) = 0.$$

Note that the local attractors are ordered by inclusion, differently from the obtained Morse sets, which are disjoint[9,19–23].

**Definition.** The unstable set of an invariant set $\Xi$ is defined by

$$W^{\mathrm{u}}(\Xi) = \{z \in X : \text{ there is a global solution } \xi : \mathbb{R} \to X \text{ for } S(t) \text{ satisfying } \xi(0) = z \text{ and such that } \lim_{t \to -\infty} \mathrm{dist}(\xi(t), \Xi) = 0\}.$$

*Remark.* Reminding that $\Xi$ is invariant it is easy to show that $\Xi \subset W^{\mathrm{u}}(\Xi)$.

**Definition.** The stable set of an invariant set $\Xi$ is defined by

$$W^{\mathrm{s}}(\Xi) = \{z \in X : \text{ such that } \lim_{t \to +\infty} \mathrm{dist}(S(t)z, \Xi) = 0\}.$$

With a Morse decomposition $\Omega = \{\Xi_1, ..., \Xi_N\}$ we can construct a sequence of local attractors setting

$$A_i = \Xi_i \cup \left[ \bigcup_{j=1}^{i-1} W^u(\Xi_j) \right].$$

And the $\mathscr{GA}$ $\mathscr{A}$ can be written as the union of the unstable manifolds related to each set in $\Omega$, i.e,

$$\mathscr{A} = \bigcup_{j=1}^{N} W^u(\Xi_j). \tag{2.1}$$

When $\Xi_j$ are equilibria $u^*_j$, the attractor is described as the union of the unstable manifolds associated to them

$$\mathscr{A} = \bigcup_{j=1}^{N} W^u(u^*_j).$$

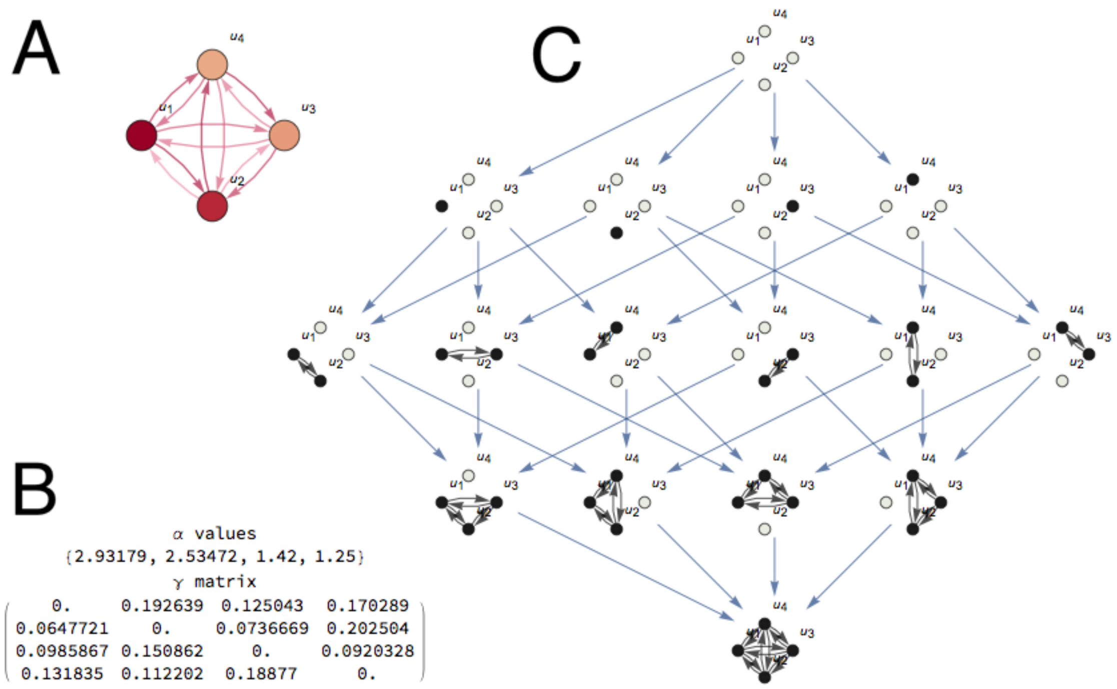


**Figure 2.2.** **A**, Structural network of a 4-dimensional ($n = 4$) Lotka-Volterra (LV) system given by (2.4). **B**, $\alpha$ and $g\Gamma$ values in this example. **C**, *Informational Structure* (IS): the global attractor $\mathscr{GA}$ is isomorphic to a directed graph. Each of the stationary points is associated with a vertex or node from the graph, and there is a directed edge from the vertex associated to one stationary point A towards the vertex associated to the stationary point B if and only if there is a global solution that connects A to B. That edge is directed because there is a Lyapunov function which is decreasing across the global solution that connects A to B. In this example the IS is a new network made by sixteen nodes ($2^n$, the maximum possible) associated to the sixteen stationary points and directed links associated to bounded solutions. The relation induced by the links is transitive in such a way that only the minimal links to understand the connecticity between nodes are represented. Each node of the IS can be represented as a subgraph of the original LV system where non-null components of the associated stationary point are shown in black and null in grey. There are 5 energy levels.

**Definition.** We will say that a semigroup $\{S(t) : t \geqslant 0\}$ with a $\mathscr{GA}$ $\mathscr{A}$ and a disjoint family of isolated invariant sets $\Omega = \{\Xi_1, ..., \Xi_N\}$ is a gradient semigroup with respect to $\Omega$ , if there exists a continuous function $V : X \to \mathbb{R}$ such that $[0, \infty) \ni t \mapsto V(S(t)u) \in \mathbb{R}$ is decreasing for each $u \in X$ , $V$ is constant in $\Xi_i$ for each $1 \leqslant i \leqslant N$ , and $V(S(t)u) = V(u)$ for all $t \geq 0$ if and only if $u \in \bigcup_{j=1}^{N} \Xi_j$.

$V$ is called Lyapunov function respect to $\Omega$.

So, a DS is called gradient if there is some continuous real-valued function which is strictly decreasing on nonconstant solutions.

It has been also proved in[16] that a semigroup $\{S(t) : t \geqslant 0\}$ is a gradient semigroup with respect to $\Omega$ if and only if it is a gradient-like semigroup with respect to $\Omega$. Essentially, this important result says that, given a disjoint family of isolated invariant sets $\Omega = \{\Xi_1, ..., \Xi_N\}$ for a semigroup $S(t)$, the dynamical property of being gradient-like, the existence of an associated ordered family of local attractor-repellers, and the existence of a Lyapunov functional related to $\Omega$, are equivalent properties.

This description shows a geometrical picture of the $\mathscr{GA}$ of a gradient-like system, in which all the stationary points or isolated invariant sets (Morse sets) are ordered by connections related to its *level of attraction*[24] or stability. Thus a consequence of it is that the $\mathscr{GA}$ is isomorphic to a directed graph: each of the $\Xi_i$ is associated with a vertex or node from the graph, and there is a directed edge from the vertex associated to $\Xi_i$ towards the vertex associated to $\Xi_j$ if and only if there is a global solution that connects $\Xi_i$ to $\Xi_j$. That edge is directed because there is a Lyapunov function which is decreasing across the global solution that connects $\Xi_i$ to $\Xi_j$. The resulting directed graph is called *Informational Structure* (IS)[25,26].

##### *2.2.3 Energy levels*

Any Morse decomposition $\Omega = \{\Xi_1, ..., \Xi_N\}$ of a compact invariant set $\mathscr{A}$ leads to a partial order among the isolated invariant sets $\Xi_i$; that is, we can define an order between two isolated invariant sets $\Xi_i$ and $\Xi_j$ saying that $\Xi_i$ preceds $\Xi_j$ ($\Xi_i \prec \Xi_j$ ) if there is a chain of global solutions

$$\{\xi_\ell,\ i \le \ell \le j-1\} \tag{2.2}$$

with

$$\lim_{t\to-\infty} dist(\xi_\ell(t), \Xi_\ell) = 0$$

and

$$\lim_{t\to\infty} dist(\xi_\ell(t), \Xi_{\ell+1}) = 0.$$

This implies that, given any dynamically gradient semigroup with respect to the disjoint family of isolated invariant sets $\Omega = \{\Xi_1, ..., \Xi_N\}$, there exists a partial order in $\Omega$ .

There exists a dynamical description of a generalized gradient-like semigroup by reordering and regrouping the corresponding isolated invariant subsets to obtain a totally ordered family of isolated invariant sets that we refer to as energy levels. This new family of isolated invariant sets is a Morse decomposition of $\mathscr{A}$ with fewer invariant sets but in such a way that it still gives us a Lyapunov function that is constant only in the solutions lying in the original isolated invariant sets. In a certain sense, this decomposition is the coarsest decomposition which still gives us a Lyapunov function which is constant only in the solutions lying in the original isolated invariant sets. In[24] (see also[22]) it is shown that there exists another Morse decomposition given by the so-called energy levels $\mathscr{N} = \{\mathscr{N}_1, \mathscr{N}_2, \cdots, \mathscr{N}_q\}$, $q \le N$.

Let us consider

$$\mathscr{M}_1 := \{\Xi_l \in \Omega :\ \text{there is no element}\ \Xi \in \Omega\ \text{that preceeds}\ \Xi_l\}$$

and, for any integer $k \ge 2$,

$$\mathscr{M}_k := \{\Xi_l \in \Omega\text{: if}\ \Xi \in \Omega\ \text{and}\ \Xi \prec \Xi_l\ \text{then}\ \Xi \in \mathscr{M}_{k-1}\}.$$

Note that, by definition, $\mathscr{M}_k \subset \mathscr{M}_{k+1}$.

We now define the sets

$$\mathscr{N}_1 := \bigcup_{\Xi \in \mathscr{M}_1} \Xi$$

and

$$\mathscr{N}_k := \bigcup_{\Xi \in \mathscr{M}_k \setminus \mathscr{M}_{k-1}} \Xi\ ,$$

for all $k \ge 2$.

Each of the levels $\mathscr{N}_i$, $1 \le i \le q$ is made of a finite union of the isolated invariant sets in $\Omega$ and $\mathscr{N}$ is totally ordered by the dynamics defined by (2.2). The total amount of levels energy, $q$, is a very important parameter in our data analysis (see Results). Indeed, the associated Lyapunov function has strictly decreasing values in any global solution linking two different level-sets of $\mathscr{N}$ and any two elements of $\Omega$ which are contained in the same element of $\mathscr{N}$ (same energy level) are not connected.

##### *2.2.4 Dynamical informational structures*

It is evident that brain dynamics does not converge or stabilize around a fixed set of invariants, but it can be described as a continuous flow of quick and irregular oscillations[27,28]. An alternative consists in the introduction of Dynamical Informational Structures (DISs), i.e., a continuous time series of ISs that determines the dynamics in each time value. We can make parameters depend on time, so that for each $t \in \mathbb{R}^+$ we have an associated IS, $\mathscr{I}_t$. Thus, the DISs ([25,26]) is defined as a continuous flow

$$\mathscr{D} : \mathbb{R}^+ \to \mathscr{G}$$

$$\mathscr{D}(t) = \mathscr{I}_t \in \mathscr{G},\ \text{for all}\ t \in \mathbb{R}^+$$

where $\mathscr{G}$ is the set of all possible graphs. Note that $\mathscr{D}$ induces a continuous movement of structures.

#### 2.3 Measuring energy levels from fMRI data

##### 2.3.1 The Lotka-Volterra model

In general, the mathematical way to describe and characterize dynamics is by (ordinary or partial) differential equations (continuous time)[29] or difference equations (discrete time)[30]. Here we choose the first one. Furthermore, many of real phenomena can be described by a set of key nodes and their associated connections, building a network[31,32]. In this way, we can simulate a real situation using an abstract graph describing its underlying architecture. Let's suppose[33] that each node $i$ is associated with a single variable $u_i$. We use $u$ to denote the vector of variables. Consider the continuous DS

$$\frac{du_i}{dt} = f_i(u_i) + \sum_{j=1}^{n} A_{ij} h_{ij}(u_i, u_j),\ i \in \{1, ..., n\}, \tag{2.3}$$

for a general model for $n$ nodes where $A = [A_{ij}]$ is the network adjacency matrix and $h_{ij}(u_i, u_j)$ gives the effect of network neighbors on each others' dynamics.

A particular case of that is the Lotka-Volterra (LV) model which has been used to generate reproducible transient sequences in neural circuits[34–38]. In our case, for a general model for $n$ nodes, we define a system of $n$ differential equations given by:

$$\frac{du_i}{dt} = u_i \left( \alpha_i - u_i + g \sum_{j=1}^{n} \gamma_{ij} u_j \right), \quad i = 1, ..., n, \tag{2.4}$$

where the matrix $\Gamma = (\gamma_{ij}) \in R^{n\times n}$ is referred to the interaction-matrix and $g \in \mathbb{R}^+$ is a parameter that modulates the coupling strength between the different nodes of the network. In matrix formulation, (2.4) reads as

$$\frac{du}{dt} = u\left(\alpha - u + g\Gamma u\right), \tag{2.5}$$

with $\Gamma \in \mathbb{R}^{n^2}$ and $\alpha \in \mathbb{R}^n$ where the product and the identity are assumed component by component. Originally LV model comes from populations dynamics and its solutions are restricted to positive values, so the phase space for (2.5) is the positive orthant

$$\mathbb{R}^n_+ = \{u = (u_1, \ldots, u_n) \in \mathbb{R}^n, u_i \geq 0, \quad i = 1, \ldots, n\}. \tag{2.6}$$

The empirical interaction matrix $\Gamma$ used in our study has diagonal entries equal zero, that is, it does not consider the interaction of a brain area with itself. To guarantee the global stability of the system, it is necessary to include the so-called logistic term $-u_i$ in (2.4) or $-u$ in (2.5) (which is equivalent to making $g\Gamma$ has a main diagonal $-1$) to guarantee (given $\Gamma$ and under certain conditions on the parameter g) the global stability of the system (see section 2.3.5).

Given an initial data for (2.4), sufficient conditions for the existence and uniqueness of global solutions are well-known (see, for instance,[39,40]). The following theorem ensures the existence of a positive solution if $g$ is smaller than a value given in the result.

**Theorem.** *Let be $u_{i0}$ with $i = 1, ..., n$ a vector of positive components and $g < 1/\rho(A)$, where $\rho(A)$ is the spectral radius of $A$. Then, the system (2.4) has a unique solution $u_i(t)$ with $i = 1, ..., n$ in any time $t > 0$ with $u_i(t) = u_{i0}$ in $t = 0$ and besides, it is positive.*

*Remark.* The spectral radius of a matrix $A$, denoted by $\rho(A)$, is the modulus of the eigenvalue with the biggest modulus; $\rho(A) = max_\lambda |\lambda(A)|$, with $\lambda(A)$ eigenvalue of $A$.

The set of equations (2.4) defines a dynamics on a structural graph with $N$ nodes, taking one equation for the description of the dynamics on each of the nodes. As we have stated before, we model the brain dynamics using LV equations since the associated ISs and its dependence on parameters is very well understood.

**Theorem.** *Each stationary point $u^* = (u^*_1, u^*_2, ..., u^*_m)$ of (2.4) consists of a unique combination of null and non-zero variables.*

*Proof.* We have to prove that given $I$ a subset of $M = \{1, ..., m\}$ such that $u^*_l = 0$ if and only if $l \in I$ the solution $u^*$ is unequivocally determined. Each stationary solution holds $u^*_i \left( \alpha_i - u^*_i + g \sum_{j=1}^{n} \gamma_{ij} u^*_j \right) = 0$ for $i = 1, ..., n$, so for each $i$ it can either hold $u^*_i = 0$ or

$$\alpha_i - u^*_i + g \sum_{j=1}^{n} \gamma_{ij} u^*_j = 0. \tag{2.7}$$

For any $k \notin I$ and due to (2.7) we have

$$u_k^* = \alpha_k + g \sum_{j \notin I}^{n} \gamma_{kj} u_j^*$$

which univocally defines a positive number since it can not be zero by definition of $I$ and can not be negative because $u^* \in \mathbb{R}_+^n$. Therefore the solution $u^*$ is univocally defined by $I$. □

Recall that as the DS is defined on a graph, two different graphs are defined: **A** the proper graph on which the DS is defined (structural network) and **B** the IS. In the particular case of the LV model there is also an unique subgraph of the structural graph associated to each stationary point (nodes of B) since each stationary point has a unique combination of null and non-zero variables and the corresponding subgraph would be the one that includes the nodes corresponding to the non-null variables and does not include any other node (we could say that it is an attracting subgraph, see Fig.2.2). Thus, the $\mathscr{G}\mathscr{A}$ and its corresponding directed graph, the IS, can be understood as *a new dynamical network describing all the possible feasible future networks*[41,42].

##### 2.3.2 Lotka-Volterra Transform

As noted, we also want each IS to flow in time. In the particular case of the dynamic system associated with the Lotka-Volterra equations we can specify the definition of DIS as follows:

$$\mathscr{D} : \mathbb{R}^+ \to 2^{\mathscr{I}}$$

$$\mathscr{D}(t) = \mathscr{I}_t \in 2^{\mathscr{I}}, \text{ for all } t \in \mathbb{R}^+$$

where $2^{\mathscr{I}}$ means the set of all subgraphs of $\mathscr{I}$ and $\mathscr{I}$ is the complete graph with all possible stationary solutions that are, at most $2^n$ according to the theorem of section 2.3.1. Thus, we can consider the LV system of differential equations driven by time dependent sources $\alpha(t)$ (and/or $\gamma_{ij}(t)$, but from now on we will consider $\gamma_{ij}$ gamma parameters not dependent on time because they represent the structural connectivity of human brain), given by:

$$\alpha : \mathbb{R}^+ \to \mathbb{R}^n$$

$$\alpha(t) = (\alpha_1(t), ..., \alpha_i(t), ..., \alpha_n(t))^T, \text{ for all } t \in \mathbb{R}^+$$

That is to say:

$$\dot{u}_i = u_i \left( \alpha_i(t) - u_i + g \sum_{j=1}^{n} \gamma_{ij} u_j \right), \quad i = 1, ..., n. \tag{2.8}$$

**Definition.** Given the values of the parameters $g \in \mathbb{R}^+$ and $\Gamma = (\gamma_{ij}) \in \mathbb{R}^{n \times n}$ in a LV model (2.8), and given a function of time $u$:

$$u : \mathbb{R}^+ \to \mathbb{R}_+^n$$

$$u(t) = (u_1(t), ..., u_i(t), ..., u_n(t))^T, \text{ for all } t \in \mathbb{R}^+$$

$$u_i(t) \in \mathbb{R}^+, \text{ for all } i = 1, ..., n.$$

we define the Lotka-Volterra transform (LVT) of $u(t)$ as the function $\alpha(t)$ that fulfills:

$$\alpha_i(t) = \frac{\dot{u}_i(t)}{u_i(t)} + u_i(t) - g \sum_{j=1}^{n} \gamma_{ij} u_j(t), \quad i = 1, ..., n.$$

*Remark.* In the previous expression we have solved for $\alpha_i(t)$ which makes sense when we have experimental data of the $u_i$, and therefore also of the $\dot{u}_i(t)$. Usually $u(t)$ is the solution of the equation given $\alpha(t)$ but now we are in the opposite situation. LVT is defined in such a way that given the initial empirical values of $u(t)$ the LV equations would fit exactly all the empirical $u(t)$ data. Nevertheless it is very important noticing that we are not assessing the LV equations as a model (it is easy to see that choosing the suitable $\alpha(t)$ the fitting is always perfect) but looking for the corresponding DISs to study them.

In practice both $u_i(t)$ and $\dot{u}_i(t)$ will be empirical values from BOLD signals in fMRI that are measured as time series, that is, we know their values for discrete values of time $t = t_k = k{\cdot}\Delta t$ where $\Delta t$ is the temporal step between meausements (in our data $\Delta t = 2.08$ seconds) and $k = 1, ..., T$ where $T$ is the length of a given data time series. We take the simplest discrete version of the equation (2.8):

$$\frac{u_{i,2} - u_{i,1}}{\Delta t} = u_{i,1}\left(\alpha_{i,1} - u_{i,1} + g\sum_{j=1}^{n}\gamma_{ij}u_{j,1}\right), \quad i = 1, ..., n \tag{2.9}$$

$$\frac{u_{i,k+1} - u_{i,k-1}}{2{\cdot}\Delta t} = u_{i,k}\left(\alpha_{i,k} - u_{i,k} + g\sum_{j=1}^{n}\gamma_{ij}u_{j,k}\right), \quad i = 1, ..., n \;\; k = 2, ..., T-1 \tag{2.10}$$

$$\frac{u_{i,T} - u_{i,T-1}}{\Delta t} = u_{i,T}\left(\alpha_{i,T} - u_{i,T} + g\sum_{j=1}^{n}\gamma_{ij}u_{j,T}\right), \quad i = 1, ..., n \tag{2.11}$$

and, solving for $\alpha$, the correspondig LVT:

$$\alpha_{i,1} = \frac{u_{i,2} - u_{i,1}}{\Delta t{\cdot}u_{i,1}} + u_{i,1} - g\sum_{j=1}^{n}\gamma_{ij}u_{j,1}, \quad i = 1, ..., n$$

$$\alpha_{i,k} = \frac{u_{i,k+1} - u_{i,k-1}}{2\Delta t{\cdot}u_{i,k}} + u_{i,k} - g\sum_{j=1}^{n}\gamma_{ij}u_{j,k}, \quad i = 1, ..., n \;\; k = 2, ..., T-1 \tag{2.12}$$

$$\alpha_{i,T} = \frac{u_{i,T} - u_{i,T-1}}{\Delta t{\cdot}u_{i,T}} + u_{i,T} - g\sum_{j=1}^{n}\gamma_{ij}u_{j,T}, \quad i = 1, ..., n$$

where $\alpha_{i,k}$ with $i = 1, ..., n$ and $k = 1, ..., T$ would be the one which actually builds the empirical BOLD signal in the following approximate solution of the LV equations:

$$u_{i,2} = u_{i,1} + \Delta t{\cdot}u_{i,1}\left(\alpha_{i,1} - u_{i,1} + g\sum_{j=1}^{n}\gamma_{ij}u_{j,1}\right), \quad i = 1, ..., n$$

$$u_{i,k} = u_{i,k-2} + 2{\cdot}\Delta t{\cdot}u_{i,k-1}\left(\alpha_{i,k-1} - u_{i,k-1} + g\sum_{j=1}^{n}\gamma_{ij}u_{j,k-1}\right), \quad i = 1, ..., n \;\; k = 3, ..., T \tag{2.13}$$

knowing the empirical $u_{i,1}$ for $i = 1, ..., n$ as initial conditions.

$n$ is the number of Regions of Interest (ROIs) in which we divide the human brain and every different $i$ from 1 to $n$ corresponds to a different ROI. At each time step $k$ there is a $\alpha_{i,k}$ column with $n$ components, so, finally we will obtain a temporal series of $T$ different ISs.

It is simple to demostrate that in certain conditions, instead of (2.13), an Explicit Runge-Kutta Method (ERKM) can be used to define the LVT. Recall that an ERKM solves $\dot{u} = f(t, u)$ by

$$u_{k+1} = u_k + \Delta t\sum_{l=1}^{s} b_l K_l$$

where

$$K_l = f(t_k + c_l\Delta t,\, u_k + \Delta t(a_{l1}K_1 + a_{l2}K_2 + ... + a_{l,l-1}K_{l-1})) \;\; l = 1, 2, ..., , s$$

specifying the integer $s$ (number of stages), the coefficients $a_{ij}$ (for $1 \le j < i \le s$ called the Runge–Kutta matrix), $b_i$ (for $i = 1, 2, ..., s$ known as the weights) and $c_i$ (for $i = 2, 3, ..., s$ known as the nodes).

**Theorem.** *An ERKM can be used to define the LVT if $c_i \in \{0, 1\}$ for $i = 1, 2, ..., s$.*

For example, is possible to define the LVT to rebuild an empirical signal using the Heuns's method, that is to say, the second-order Runge-Kutta method with two stages, $c_1 = 0$ and $c_2 = 1$.

It should not be surprising that conditions on an ERKM have to be imposed to be able to use it in the definition of the LVT because they both try to solve different problems. An ERKM is used when we know the explicit dependence of the differential equation in time which in our case is $\alpha(t)$ and we try to find out the solution of the equation $u(t)$. The LVT is used when we know the solution to the equation $u(t)$ and try to find out $\alpha(t)$.

##### 2.3.3 Linear transform of BOLD signal

The fMRI technique relies on the fact that cerebral blood flow and neuronal activation are coupled. In this work we consider the BOLD signal as an indicator of activity in each brain area, but before assessing alpha as the LVT we calculate:

-First, the z-scores of the BOLD signal:

$$z = \frac{x-\mu}{\sigma}$$

where $x$ is the BOLD signal, $\mu$ is the mean of the sample and $\sigma$ is the standard deviation of all samples.

-Then we add a constant b:

$$\bar{u} = z + b$$

choosing $b$ in order to obtain $\bar{u} > 0$ since due to biological reasons LV equations do not make sense for negative values and the LVT is undefined for $\bar{u} = 0$.

-Finally, since the LV equations we use in this work are non-dimensional, before empirical data are substituted in them we must determine a characteristical $t_c$ and $u_c$ and substitute the non-dimensional values $t = \bar{t}/t_c$ and $u = \bar{u}/u_c$ (where $\bar{t}$ and $\bar{u}$ are dimensional values) in the non-dimensional LV equations:

$$\frac{du}{dt} = u(\alpha - u + g\Gamma u) \Rightarrow \frac{t_c}{u_c}\frac{d\bar{u}}{d\bar{t}} = \frac{1}{u_c^2}\bar{u}(\bar{\alpha} - \bar{u} + g\Gamma\bar{u}) \Rightarrow u_c t_c \frac{d\bar{u}}{d\bar{t}} = \bar{u}(\bar{\alpha} - \bar{u} + g\Gamma\bar{u})$$

$u_c t_c$ is the same for each component $i = 1,...,N$ and for asleep and awake subjects but determined optimizing the paired difference test between both brain states.

##### 2.3.4 Lineal Complementary Problem

###### Some preliminary concepts

**Definition.** We call $X$ in $\mathbb{R}^n$ a cone if, for any $x \in X$ and $t \geq 0$, $tx \in X$ holds. If $X$ is convex, it is called a convex cone.

**Definition.** Given $B \in \mathbb{R}^{m\times p}$, we define $pos(B)$ as

$$pos(B) = \left\{q \in \mathbb{R}^m : q = Bv \text{ for } v \in \mathbb{R}^p_+\right\}.$$

Columns of $B$, $B_{.i}$, for $i = 1,...p$, are defined as the generators of **pos(B)**.

**Definition.** Column vectors of $M \in \mathbb{R}^{n\times n}$, written as $M_{.j}$, and columns of identity matrix $I \in \mathbb{R}^{n\times n}$ written as $I_{.j}$, generate the complementary pair of column vectors $\left\{I_{.j}, -M_{.j}\right\}$ for $j = 1,...,n$. If $B_{.j} \in \left\{I_{.j}, -M_{.j}\right\}$, for $j = 1,...,n$, the set $(B_{.1},...,B_{.n})$ is defined as a complementary vector set.

**Definition.** Let $M$ of order $n \times n$ and $(B_{.1},...,B_{.n})$ a complementary vector set.

$$pos(B_{.1},...,B_{.n}) = \{y : y = \alpha_1 B_{.1} + ... + \alpha_n B_{.n};\ \alpha_i \geq 0\}$$

is called a complementary cone.

$\mathscr{C}(M)$ is the set of all complementary cones associated with $M$. Given $M$, there exist $2^n$ complementary cones in $\mathscr{C}(M)$ since each column $B_{.j}$ has two possibilities: $I_{.j}$ or $-M_{.j}$.

**Lineal Complementary Problem** The Linear Complementary Problem (LCP) (see[43,44]) states that, given $r \in \mathbb{R}^n$ and a matrix $M$ of order $n$, we try to find $(w,z) \in \mathbb{R}^{2n}$ , $w = (w_1, w_2, ..., w_n)^T$ , $z = (z_1, ..., z_n)^T$ , such that

$$w = r + Mz$$
$$w \geq 0,\ z \geq 0 \text{ and } w_i z_i = 0 \text{ for all } i = 1,...,n.$$

Observe that the $LCP(r,M)$ is equivalent to find a cone in $\mathscr{C}(M)$ containing $q$, i.e., to get $(w,z) \in \mathbb{R}^{2n}$ satisfying

$$\begin{aligned} Iw - Mz &= r \\ w,z &\geq 0 \\ w_j z_j &= 0, \text{ for } j = 1,\cdots,n. \end{aligned} \tag{2.14}$$

Thus, $w_j$ is associated with column vector $I_{.j}$ and $z_j$ to $-M_{.j}$ (see[44,45]).

Existence and uniqueness of solution to the LCP depends on the stability of the matrix $M$.

**Definition.** $A \in \mathbb{R}^{n\times n}$ is said to be stable if all associated eigenvalues has negative real part.

**Definition.** $A$ is positive semi-definite (negative semi-definite), if $u^T Au \geq 0$ ($u^T Au \leq 0$) for all $u \in \mathbb{R}^n$. It is positive definite (negative definite) if $u^T Au > 0$ ($u^T Au < 0$) for all $u \in \mathbb{R}^n \setminus \{0\}$.

a) $A$ belongs to class $S_w$ or is Lyapunov-stable (see[46]), $A \in S_w$, if there exists a diagonal positive matrix $W$ such that $WA + A^T W$ is negative definite.

b) $A$ is called negative dominant diagonal, $A \in NDD$ if there exist $n$ positive numbers $v_i > 0$ such that

$$-v_i a_{ii} > \sum_{i\neq j}^{n} |a_{ij}| v_j, \quad i = 1, \ldots, n.$$

c) Recall that a minor of a matrix $A$ is the determinant of some smaller square matrix, cut down from $A$ by removing one or more of its rows or columns. If $I$ and $J$ are subsets of $\{1,...,n\}$ with k elements, then we write $[A]_{I,J}$ for the $k\times k$ minor of $A$ that corresponds to the rows with index in $I$ and the columns with index in $J$. If $I = J$, then $[A]_{I,J}$ is called a principal minor. $A$ is said to be a $P$-matrix, $A \in P$, if all principal minors of $A$ are positive.

A matrix $A$ in $S_w$ implies $-A$ to be a $P-$ matrix ([40]).

Sufficient conditions for $A$ to be in $S_w$ are shown in the following result (see[47]): $A$ belongs to $S_w$ if any of the following conditions hold:

(i) $A$ is negative diagonal dominant;

(ii) $A$ es negative definite.

Next result (see[44]) gives existence and uniqueness of solution for the (LCP).

**Theorem.** *The $LCP(r,M)$ possesses a unique solution $r \in \mathbb{R}^n$ if and only if $M$ is a $P-matrix$.*

The following results are important in order to see the continuous dependence between $r$ of the $LCP(r,M)$ and its associated solution, which will be crucial to observe the dynamical behaviour of ISs.

**Definition.** A class of convex cones in $\mathbb{R}^n$ is a partition of $\mathbb{R}^n$ if
Each cone has a nonvoid interior.
The union of all cones is $\mathbb{R}^n$.
Each pair of the interior of cones is disjoint.

**Theorem.** *Let $M$ a matrix of order $n$. The class of complementary cones $\mathscr{C}(M)$ is a partition of $\mathbb{R}^n$ if and only if $M$ is a $P$-matrix.*

**Definition.** A function $f : U \subset \mathbb{R}^n \mapsto \mathbb{R}^m$ is linear by parts, if $f$ is continuous and domain $U$ is the union of a finite number of convex polyhedra $P_i$, and $f$ is linear on each $P_i$.

**Theorem.** *Suppose that the LCP($r,M$) has a unique solution $z$. Then the function $L : \mathbb{R}^n \to \mathbb{R}^n$ defined as $L(r) = z$ is linear by parts (see[44]).*

##### 2.3.5 LV systems and the LCP

Consider the stationary point $u^* = (u_1^*, u_2^*, ..., u_n^*)$ of the LV equations expressed in the general way:

$$\frac{du_i}{dt} = u_i \left( b_i + \sum_{j=1}^{n} a_{ij} u_j \right), \quad i = 1, ..., n, \tag{2.15}$$

Recall that for us $b_i = \alpha_i$ and $a_{ij} = -\delta_{ij} + g\hat{A}\cdot\gamma_{ij}$ where $\delta_{ij}$ is the Kronecker delta. In a stationary point $\frac{du_i}{dt} = 0$ for $i = 1, ..., n$ and by (2.6) $u_i^* \geq 0$. So any stationary point $u^*$ holds that:

$$\begin{cases} u_i^* & \geq & 0, \\ u_i^* \left( b_i + \sum_{j=1}^{n} a_{ij} u_j^* \right) & = & 0, \quad i = 1, ..., n. \end{cases} \tag{2.16}$$

There exists and equivalence between looking for stationary solutions of the LV systems and the solution of a $LCP$, as shown by the following result:

**Lemma.** *The LCP$(-b,-A)$ where $b=(b_1,b_2,...,b_n)^T$ and $A=[a_{ij}]$ is equivalent to find a nonnegative $u^*$ stationary point of (2.15) satisfying:*

$$b_i+\sum_{j=1}^{n} a_{ij}u_j^* \leq 0 \quad for \;\; i=1,...,n. \tag{2.17}$$

*Proof.* It is enough to take

$$z=u^* \quad \text{y} \quad w=-b-Au^*,$$

so that $M=-A$ and $r=-b$ and thus it holds $w=Mz+r$, $w\geq 0$, $z\geq 0$ and $w_iz_i=0$, for all $i=1,\dots,n$. □

**Definition.** $u^*$ satisfying (2.17) is called saturated.

In this way, LV equations (2.15) has a unique saturated equilibrium point for each $b\in\mathbb{R}^n$ if and only if $-A$ is a $P-matrix$.

**Definition.** For a nonnegative stationary solution $u^*=(u_1^*,u_2^*,...,u_n^*)$ we define $I$ and $J$ as subsets of $N=\{1,...,n\}$ such that $u_i^*=0$ for $i\in I$ and $J=N\backslash I$.

We also define

$$\mathbb{R}_I^n=\left\{u=(u_1,u_2,...,u_n)\in\mathbb{R}_+^n \,\middle|\, u_i\geq 0 \;\text{ para } i\in I \;\text{ and } u_j>0 \text{ for } j\in J\right\}. \tag{2.18}$$

Note that $\mathbb{R}_I^n=int(\mathbb{R}_+^n)$ if $u^*>0$, i.e, if $J=N$.

The following important result (see[47]) now gives us the global stability of saturated equilibria for LV systems:

**Theorem.** *Suppose $A\in S_w$. Then the LV system (2.15) possesses a saturated stationary point $u^*$ for each $b\in\mathbb{R}^n$ which is globally stable in $\mathbb{R}_I^n$.*

It assures existence and global stability of stationary points for (2.17) (see[40]).
Note that if $A\in S_w$, every principal submatrix of $A$ also belong to $S_w$.

We finally have (see[40]):

**Corollary.** *If $A\in S_w$, then the LV system and all its associated subsystems possess a unique globally stable equilibria for each $b\in\mathbb{R}^n$.*

**Theorem.** *The system (2.5) has an asymptotically globally stable solution if $g<1/\rho(A)$.*

*Proof.* Let us write the system (2.5) $\dfrac{du}{dt}=u(\alpha-u+g\Gamma u)$ as $\dfrac{du}{dt}=u(\alpha+Au)$, where the product and the identity are asumed component by component and $A$ matrix is given by $A=-I+g\Gamma$. As we said $A\in S_w$, if there exists a diagonal positive matrix $W$ such that $WA+A^TW$ is negative definite. For these matrices there is globally stable solution (see corolllary). If $g<1/\rho(A)$, then $-1+g\lambda(A)<0$, being $\lambda(A)$ an arbitrary eigenvalue of $A$. So $B$ is negative definite, and belonging to $S_w$ trivially. □

###### *2.3.6 Summary and specification*

As a summary of the whole section and distinguishing for the case of the LV model which we are going to use, we can say that:

(i) If we suposse that $A$ in $\dfrac{du}{dt}=u(\alpha(t)+Au)$ is Lyapunov-stable there is a $\mathcal{GA}$ in the positive cone for each $t_0$ fixed, with Morse decomposition which consists of a finite set of stationary points $U^*:=\{u_1^*,u_2^*,...,u_m^*\}$ .

(ii) The semigroup associated with the equations $\dfrac{du}{dt}=u(\alpha(t_0)+Au)$ , for $t_0$ arbitrary but fixed, is a *gradient semigroup* with respect to $U^*$, so the IS is associated with that $\mathcal{GA}$ is a directed graph in such a way that there is a single point stationary globally asymptotically stable solution (GASS) in the positive cone.

(iii) This GASS can be interpreted as the lower end of the IS if it is represented with the directed links going from up to down.

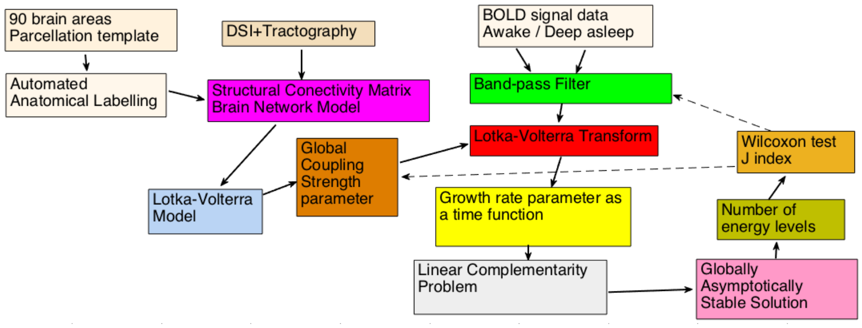


**Figure 2.3.** A more detailed flowchart than Fig. 3 illustrating the methods. First, we use the structural connectivity between 90 standardized brain areas through an automated anatomical labelling (AAL) atlas. The corresponding tractogram was obtained in a previous study[1] through diffusion tensor imaging (DTI) providing an structural connectivity network. We assume that neural activity in different brain areas can be simulated in the ansatz of interacting species in a cooperative Lotka-Volterra (LV) system constrained by the structural connectivity and a global coupling strength parameter value. With these constraints, the ansatz defines the LV transform, a mathematical operator that calculates the growth rate as a time function that reproduces exactly the filtered empirical BOLD fMRI signals. Then solving the linear complementarity problem we calculate the globally asymptotically stable stationary solution (GASS) and the number of energy levels (NoEL) of the corresponding informational structure at each time instant. We repeat this process for each subject data in resting state and in N3 state. In order to assess how different the distributions of the NoEL for awake and for deep asleep are, we calculate a statistical hypothesis test (Wilcoxon test) and the $J_{\text{ind}}$. Finally, these results are used to fit the initial values of the global coupling parameter and even the extreme values in the frequency range used for filtering.

(iv) Given $A$ Lyapunov-stable, the IS is conditioned by the GASS and the GASS is determined by what particular cone in $\mathscr{C}(-A)$ contains $-\alpha(t_0)$, which is equivalent to finding the cone in $\mathscr{C}(A)$ containing $\alpha(t_0)$.

(v) This $\alpha(t_0) \in \mathbb{R}^n$ will be in a specific cone and that determines what is the point of the attractor or single stationary point that is asymptotically stable, the GASS.

(vi) Let us remember that in a LV system $\dfrac{du}{dt} = u(\alpha(t_0) + Au)$ with $u \in \mathbb{R}^N$each stationary point has a unique combination of null and non-zero variables (see section 2.3.1). So the IS is isomorphic to a subgraph of the directed hypercube $[0,1]^N$ which has $2^N$vertices and where each directed link goes to the vertex nearest $(0,0,...,0)$ to the vertex nearest $(1,1,...1)$.

(vii) The intersections of the cones where $\alpha$ moves, and which cover all $\mathbb{R}^{90}$, are bifurcation zones. Being close to a vertex, of an edge, of a face, will indicate that with a slight change in the parameter $\alpha$ (actually, in 90 parameters, each of the components of $\alpha$), the solution is going to behave in a very different way in the future[48,49]. Note that the continuous dependence on the parameters such as on the strength of connections[50] in our characterization of the IS allows to understand the appearance of sudden bifurcations[51].

#### 2.4 Indentifying brain states: paired difference test and J index

To assess differences in the DISs between wakefulness and deep sleep, we used the non-parametric Wilcoxon signed-rank test. That is a paired difference test, tipically used when the same subject is measured before and after a treatment and that determines if two paired samples were selected from populations having the same distribution. Also, the paired Student's test can be used when the population can be assumed to be normally distributed, but we do not have enough data to know if that holds in our case so we will use the Wilcoxon signed-rank test. We consider two quantitative descriptors of the DISs, the time average of NoEL ($\overline{q}$) and the standard deviation of NoEL ($\sigma_q$), both calculated for each participant in the two brain states. We obtained the p-value of a paired and two-sided test for the null hypothesis that the distribution of average NoEL (or their standard deviation) presented the same median during wakefulness vs. deep asleep subjects. So a small p-value of that null hypothesis means that $\overline{q}$ (or $\sigma_q$) are quite different for conscious and unconscious subjects.

In adition to the p-value in the Wilcoxon test we define another measure called $J_{ind}$:

$$J_{ind} = \frac{\sum_{i=1}^{ns} (\overline{q}_{as,i} - \overline{q}_{aw,i})}{\sum_{i=1}^{ns} |\, \overline{q}_{as,i} - \overline{q}_{aw,i} \,|} \tag{2.19}$$

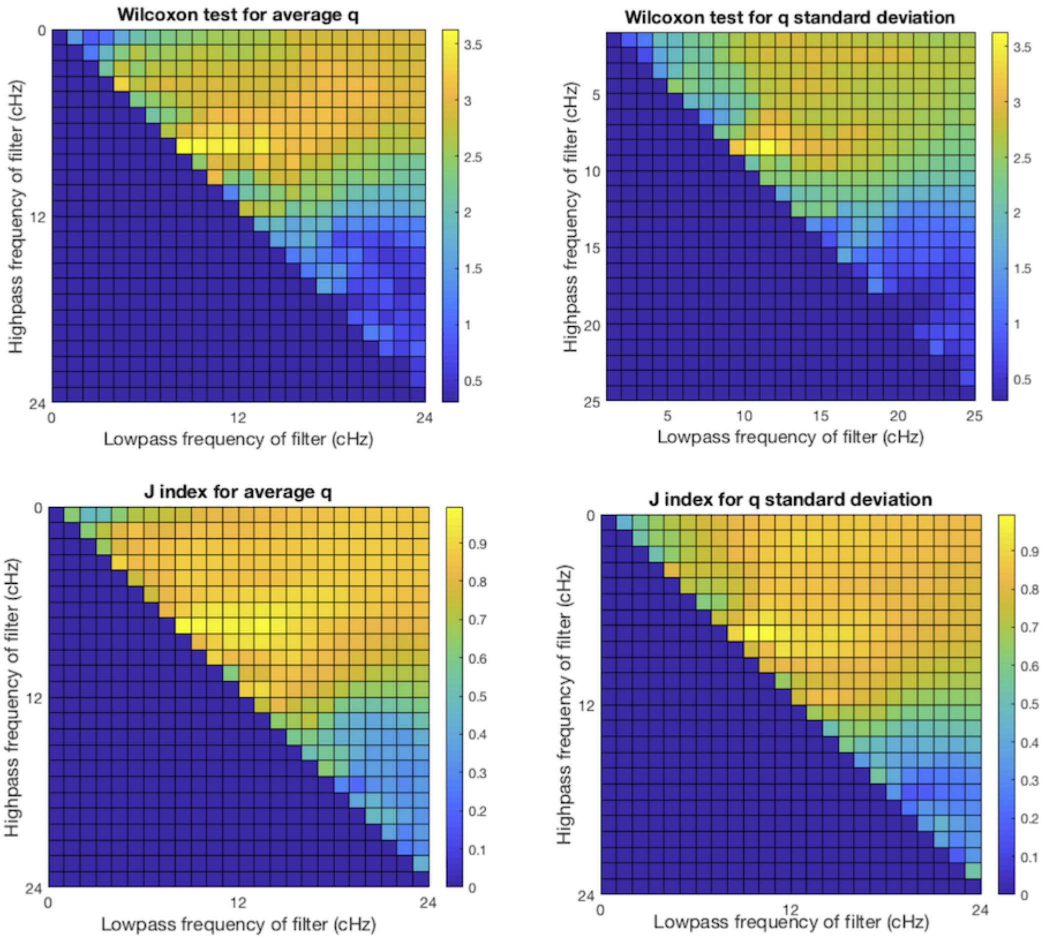


**Figure 2.4.** 2D version of Fig. 4B. Fitting of the filter used taking as an optimization criterion the minimization of the p-value of the Wilcoxon test and the $J_{\text{ind}}$ for the samples of both, $\overline{q}$ and $\sigma_q$. Since fMRI data are collected with time to repetition (TR) in the order of $2.08\,sec$ the Nyquist frequency, $0.24\,Hz$, is the frequency upper limit. The points below and to the left of the principal diagonal are meaningless since the lower end of the interval can not be greater than the upper limit and the worst value of the scale has been arbitrarily assigned to them. Both, the Wilcoxon test p-value and the $J_{\text{ind}}$ are functions of the ends of the filtering range. All values lower than $10^{-2}$ ($< 1\%$, green, yellow, orange or red) are acceptable from the point of view of hypothesis contrast. Near the principal diagonal the width of the filter is narrower so it will be easier to locate the frequencies that characterize the awake state compared to deep asleep state. For the samples of $\overline{q}$ the filter with maximum $J_{\text{ind}}$ (0.9948) was $0.08 - 0.1\,Hz$. As in the fitting of the coupling strength parameter $g$, the Wilcoxon test and the $J_{\text{ind}}$ measurements correlate, since the two graphs of each column are quite similar.

where $ns$ is the number of subjects , $\overline{q}_{as,i}$ is the value of $\overline{q}$ for subject $i$ while asleep and $\overline{q}_{aw,i}$ is the value of $\overline{q}$ for subject $i$ while awake. Then, $J_{\text{ind}} = 1$ ($J_{\text{ind}} = -1$) if the mean NoELs is larger (smaller) for deep sleep than wakefulness for all subjects and $J_{\text{ind}} \simeq 0$ if $\overline{q}$ did not depend on the brain state, that is to say, if the null hypothesis holds. $J_{\text{ind}}$ can also be defined for $\sigma_q$ or for any other parameter.

$J_{ind}$ is not really the result of a paired diffrence test but a new measure of association between a categorical variable (for example, the brain state) and a quantitative variable (for example, the mean of the NoEL). It applies to paired data sets and measures the separation between one set of data and another. It is similar to a paired case of the Cohen's $d$ index. (Cohen's $d$ is an effect size used to indicate the standardised difference between two means and can be used to accompany reporting of t-test and analysis of variance results.)

It is important not to mix up the signed-rank test with the rank sum test, both from Wilcoxon. This last test assumes that the two samples are independent while the first, which we are using here, is for paired samples.

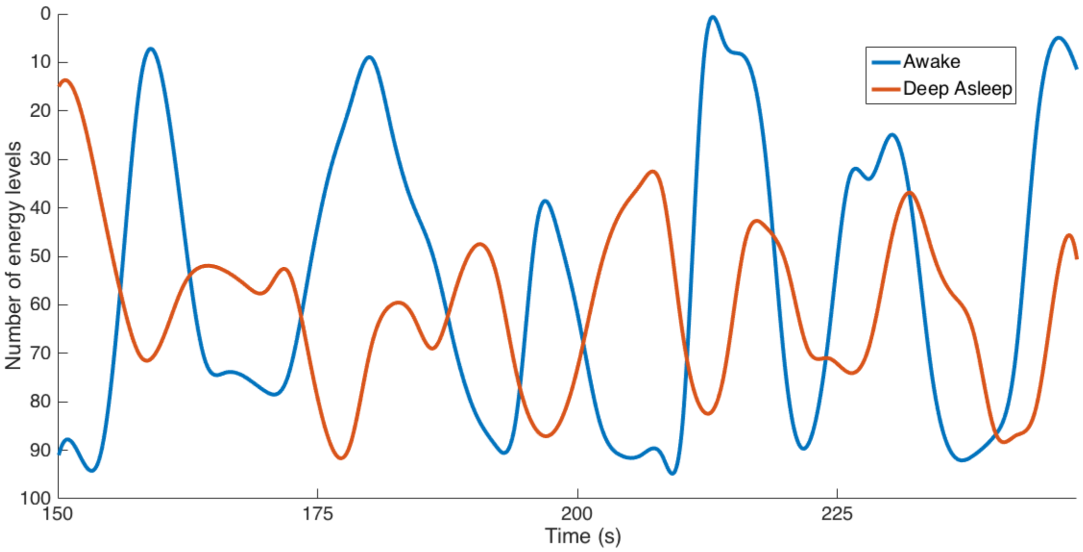


**Figure 2.5.** Similar graphic than Fig. 5 but using a different filter. Again subjects in awake condition show greater variability than for deep asleep condition. In this graphic the Y-axis is inverted: as it is said in Discussion if we use as a reference the energy of the unstable trivial solution $(0, 0, ..., 0)$ and take into account that when the number of levels increases, the stability of the GASS grows and its energy decreases, the average energy of the deep asleep state is less than the awake one. So, although the average number of energy levels (NoEL) is higher for deep asleep than for awake condition again the average energy in awake state is higher. In this100-seconds long sample of the NoEL for a different subject in both states empirical data was band-pass filtered in the range $0.04-0.07$ Hz (in Fig. 5 the range was $0.077-0.096$ Hz). Again $g=0.29$. In practice both the BOLD signal data and the NoEL are evaluated as time series, that is, for discrete values of time ($\Delta t = 2.08$ seconds). Only for this figure the discrete data were interpolated using cubic spline.

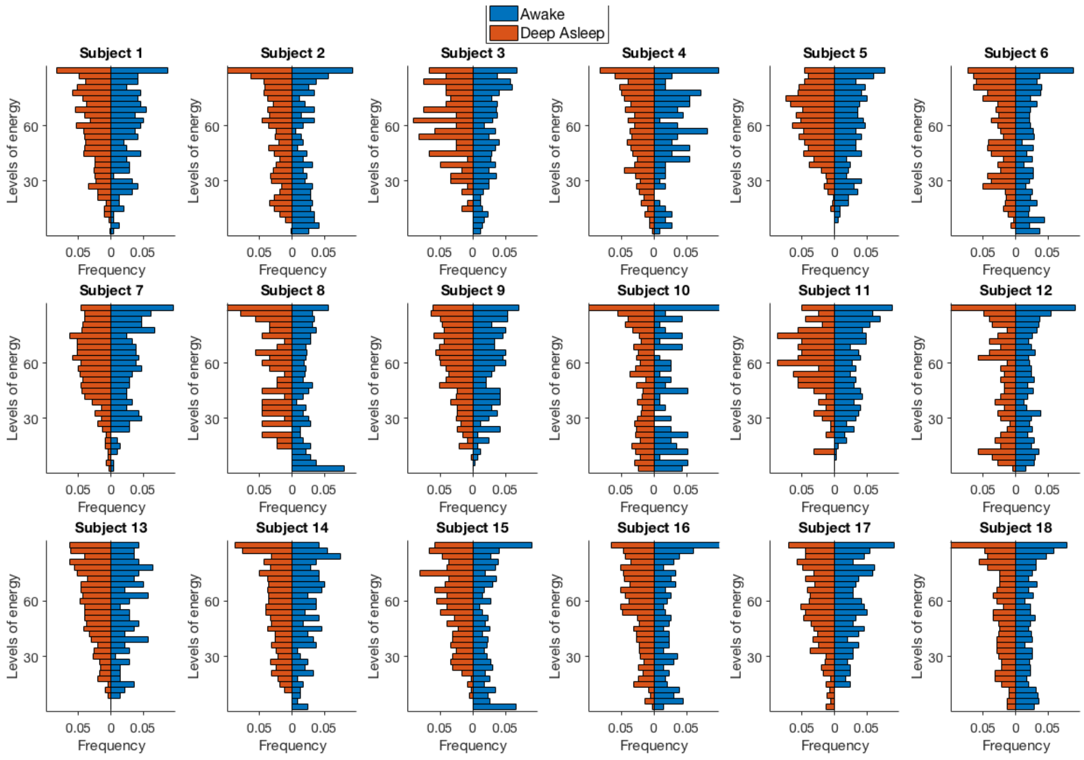


**Figure 2.6.** It is the same plot than Fig. 6B but non-smoothed back to back histograms of the distributions of the NoEL for 18 healthy subjects in awake and in deep asleep states. Each bar comprises 3 energy levels. The NoEL was obtained from filtered data between $0.077-0.096\,Hz$ and $g = 0.29$. The pattern observed is that awake distributions are more homogeneously distributed among all the possible values of energy.

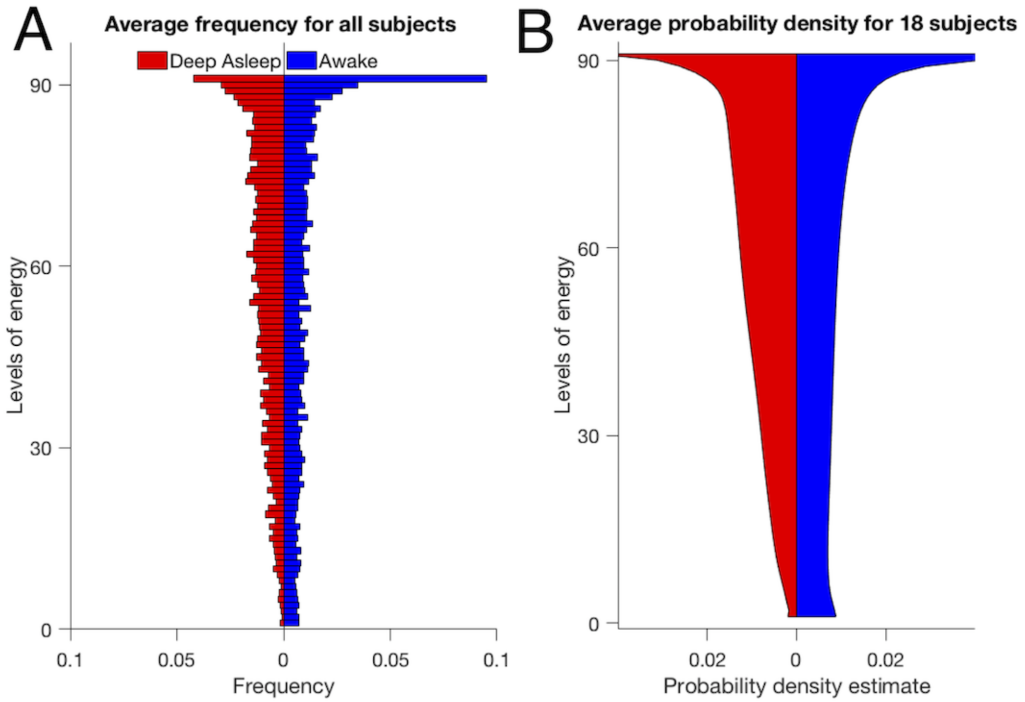


**Figure 2.7.** **A**, Non-smoothed back to back histogram of the average distribution over 18 subjects in awake and deep asleep conditions. Each bar corresponds to one energy level. Energy levels were obtained from filtered data between $0.077-0.096$ Hz and $g = 0.29$. The pattern observed is that asleep distributions tend to have higher values ($>45$) of NoEL. **B**, This Distribution estimations is the smoothed back to back histogram of the average distribution over 18 subjects in awake and deep asleep conditions. The smoothing was made using the kernel density estimation, i.e., a non-parametric way to estimate the probability density function. Inferences about the population are made based on a finite data sample. Energy levels were obtained from filtered data between $0.077-0.096$ Hz and $g = 0.29$. It is the same plot than Fig. 6C but back to back. Informational structures with low ($<28$) and very high ($>87$) numbers of energy levels are more frequent in awake subjects.

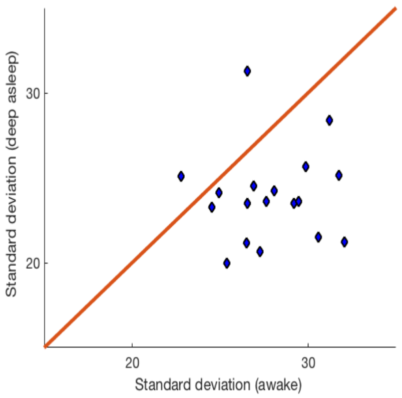


**Figure 2.8.** Standard deviation of the NoEL for deep asleep state as a function of NoEL standard deviation for awake state with filtering in the range $0.04-0.07$ Hz. The standard deviation is the same for awake and for deep asleep state along the red line. Clearly, in 16 out of 18 subjects the standard deviation is larger in awake than in deep asleep state. In Fig. 7B the filter was $0.077-0.096$ Hz and in 17 out of 18 subjects the standard deviation was larger in awake. So, the results depend on the filter but the general resolution remains the same.

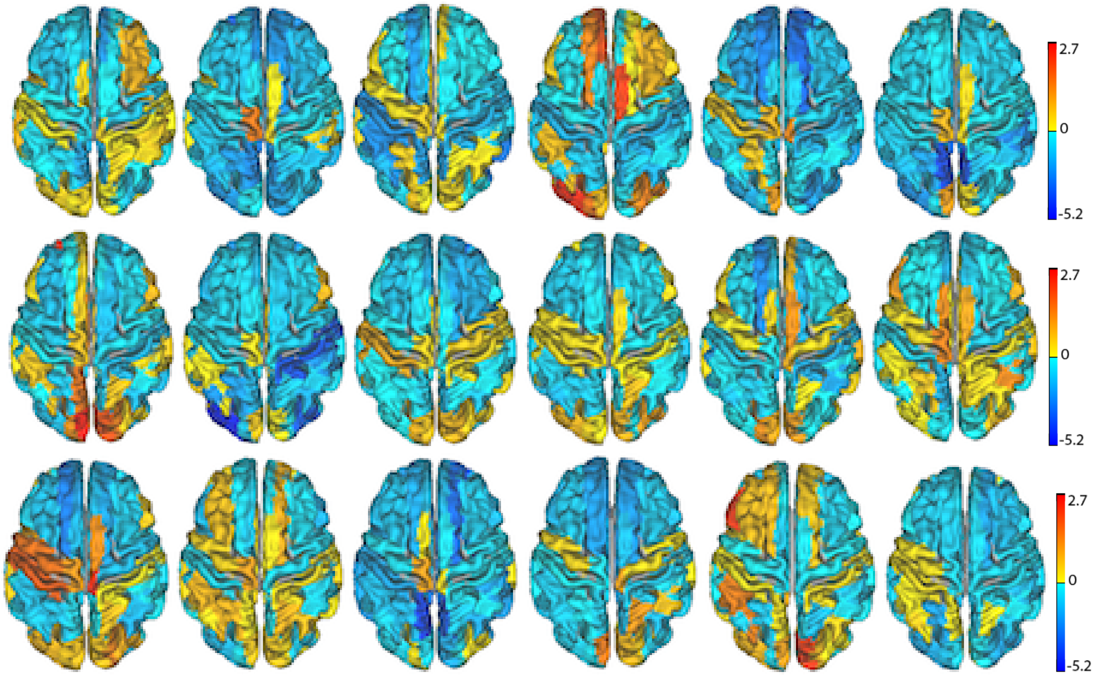


**Figure 2.9.** Difference between the frequencies with which each area appears in the GASS in awake and in deep asleep states for the 18 subjects. This difference is expressed in standard error units where the standard error is estimated by the intrapersonal variability. Given that, on average, the attractor of the sleeping condition is more populated (greater number of energy levels) than that of the awakened ones, most of the brain areas tend to appear more frequently in the attractor when subject are sleeping, i.e., most areas have cold colors.

### 3 Supplementary results

Here, we characterize the different levels of consciousness by analyzing the variability of the IS across time (see Fig.2.3 with the flow of steps applied).

First, we build a whole-brain ansatz based on the Lotka-Volterra (LV) equations for the brain dynamics, where the structural connectivity matrix obtained from empirical tractography-based DTI is chosen as the interaction matrix of the dynamical system (DS).

Then, we apply the Lotka-Volterra Transform (LVT) to calculate the growth rate $\alpha$ that reproduces exactly the filtered empirical BOLD fMRI signals.

After solving the linear complementarity problem we calculate the globally asymptotically stable solution (GASS) and the number of energy levels (NoEL) of the corresponding Informational Structure (IS) at each time instant (Fig.2.2).

In order to assess the differences between the distributions of the average NoEL ($\overline{q}$) and decide which conscious state the subject undergoes, we apply statistical tests, namely Wilcoxon and $J_{\text{ind}}$ as defined in (2.19).

### References


1. Deco, G. *et al.* Single or multiple frequency generators in on-going brain activity: A mechanistic whole-brain model of empirical meg data. *NeuroImage* **152**, 538 – 550, DOI: https://doi.org/10.1016/j.neuroimage.2017.03.023 (2017).
2. Cabral, J., Kringelbach, M. L. & Deco, G. Functional graph alterations in schizophrenia: A result from a global anatomic decoupling? *Pharmacopsychiatry* **45**, S57–S64, DOI: 10.1055/s-0032-1309001 (2012).
3. Babin, A. & Vishik, M. *Attractors of Evolution Equations*. Studies in Mathematics and its Applications (Elsevier Science, 1992).
4. Hale, J. *Asymptotic Behavior of Dissipative Systems*. Mathematical surveys and monographs (American Mathematical Society, 1988).
5. Henry, D. B. *Geometric theory of semilinear parabolic equations* (Springer-Verlag, Berlin, 1981).
6. Ladyzhenskaya, O. A. *Attractors for semigroups and evolution equations* (Cambridge University Press, 1991).

7. Robinson, J., Crighton, D. & Ablowitz, M. *Infinite-Dimensional Dynamical Systems: An Introduction to Dissipative Parabolic PDEs and the Theory of Global Attractors*. Cambridge Texts in Applied Mathematics (Cambridge University Press, 2001).

8. Temam, R. *Infinite-Dimensional Dynamical Systems in Mechanics and Physics*. Applied Mathematical Sciences (Springer New York, 1997).

9. Conley, C. *Isolated invariant sets and the Morse index*. No. 38 in CBMS Regional Conference Series in Mathematics (American Mathematical Society, Providence, 1978).

10. Joly, R. & Raugel, G. Genertic morse-smale property for the parabolic equation on the circle. *Annales de l'Institut Henri Poincare (C) Non Linear Analysis* **27**, 1397–1440, DOI: https://doi.org/10.1016/j.anihpc.2010.09.001 (2010).

11. Hale, J., Magalhaes, L. & Oliva, W. *An Introduction to Infinite Dimensional Dynamical Systems - Geometric Theory*. Applied Mathematical Sciences (Springer New York, 2013).

12. Palis, J., Manning, A. & de Melo, W. *Geometric Theory of Dynamical Systems: An Introduction* (Springer New York, 2012).

13. Ott, E. *Chaos in Dynamical Systems* (Cambridge University Press, 2002).

14. Strogatz, S. *Nonlinear Dynamics And Chaos*. Studies in nonlinearity (Sarat Book House, 2007).

15. Wiggins, S. *Introduction to Applied Nonlinear Dynamiccal Systems and Chaos*. Introduction to Applied Nonlinear Dynamiccal Systems and Chaos, (Springer-Verlag New York, I., 2003).

16. Aragao-Costa, E. R., Caraballo, T., Carvalho, A. N. & Langa, J. A. Stability of gradient semigroups under perturbations. *Nonlinearity* **24**, 2099 (2011).

17. Babin, A. V. & Vishik, M. Regular attractors of semigroups and evolution equations. *Math. Pures et Appl.* **62**, 441–491 (1983).

18. Carvalho, A., Langa, J. & Robinson, J. *Attractors for infinite-dimensional non-autonomous dynamical systems*. Applied Mathematical Sciences (Springer New York, 2012).

19. Rybakowski, K. P. *The homotopy index and partial differential equations*. Universitext (Springer-Verlag, 1987).

20. Norton, D. E. The fundamental theorem of dynamical systems. *Commentationes Math. Univ. Carol.* **36**, 585–597 (1995).

21. Hurley, M. Chain recurrence, semiflows, and gradients. *J. Dynam. Differ. Equations* **7**, 437–456, DOI: 10.1007/BF02219371 (1995).

22. Patrão, M. & San Martin, L. A. B. Semiflows on topological spaces: chain transitivity and semigroups. *J. Dynam. Differ. Equations* **19**, 155–180, DOI: 10.1007/s10884-006-9032-3 (2007).

23. Patrão, M. Morse decomposition of semiflows on topological spaces. *J. Dyn. Differ. Equations* **19**, 181–198, DOI: 10.1007/s10884-006-9033-2 (2007).

24. Aragao-Costa, E. R., Caraballo, T., Carvalho, A. N. & Langa, J. A. Continuity of lyapunov functions and of energy level for a generalized gradient semigroup. *Topol. Methods Nonlinear Anal.* **39**, 57–82 (2012).

25. Esteban, F. J., Galadí, J. A., Langa, J. A., Portillo, J. R. & Soler-Toscano, F. Informational structures: A dynamical system approach for integrated information. *PLOS Comput. Biol.* **14**, 1–33, DOI: 10.1371/journal.pcbi.1006154 (2018).

26. Kalita, P., Langa, J. A. & Soler-Toscano, F. Informational structures and informational fields as a prototype for the description of postulates of the integrated information theory. *Entropy* **21**, DOI: 10.3390/e21050493 (2019).

27. Golos, M., Jirsa, V. & Dauc´, E. Multistability in large scale models of brain activity. *PLOS Comput. Biol.* **11**, 1–32, DOI: 10.1371/journal.pcbi.1004644 (2016).

28. Deco, G. & Jirsa, V. K. Ongoing cortical activity at rest: Criticality, multistability, and ghost attractors. *J. Neurosci.* **32**, 3366–3375, DOI: 10.1523/JNEUROSCI.2523-11.2012 (2012). http://www.jneurosci.org/content/32/10/3366.full.pdf.

29. Hirsch, M., Smale, S. & Devaney, R. *Differential Equations, Dynamical Systems, and an Introduction to Chaos* (Elsevier Science, 2012).

30. Sandefur, J. *Discrete Dynamical Systems: Theory and Applications* (Clarendon Press, 1990).

31. Mortveit, H. & Reidys, C. *An Introduction to Sequential Dynamical Systems* (Springer US, 2008).

32. Osipenko, G. *Dynamical Systems, Graphs, and Algorithms*. Lecture Notes in Mathematics (Springer Berlin Heidelberg, 2006).

33. Porter, M. & Gleeson, J. *Dynamical Systems on Networks: A Tutorial*. Frontiers in Applied Dynamical Systems: Reviews and Tutorials (Springer International Publishing, 2016).

34. Afraimovich, V. S., Moses, G. & Young, T. Two-dimensional heteroclinic attractor in the generalized Lotka-Volterra system. *Nonlinearity* **29**, 1645, DOI: 10.1088/0951-7715/29/5/1645 (2016). 1509.04570.

35. Afraimovich, V. S., Zhigulin, V. P. & Rabinovich, M. I. On the origin of reproducible sequential activity in neural circuits. *Chaos: An Interdiscip. J. Nonlinear Sci.* **14**, 1123–1129, DOI: 10.1063/1.1819625 (2004). https://doi.org/10.1063/1.1819625.

36. Afraimovich, V., Tristan, I., Varona, P. & Rabinovich, M. Transient dynamics in complex systems: Heteroclinic sequences with multidimensional unstable manifolds. *Discontinuity, Nonlinearity, Complex.* **2(1)**, 21–41, DOI: 10.5890/DNC.2012.11.001 (2013).

37. Muezzinoglu, M. K., Tristan, I., Huerta, R., Afraimovich, V. S. & Rabinovich, M. I. Transients versus attractors in complex networks. *Int. J. Bifurc. Chaos* **20**, 1653–1675, DOI: 10.1142/S0218127410026745 (2010). https://doi.org/10.1142/S0218127410026745.

38. Rabinovich, M., Varona, P., Tristan, I. & Afraimovich, V. Chunking dynamics: heteroclinics in mind. *Front. Comput. Neurosci.* **8**, 22, DOI: 10.3389/fncom.2014.00022 (2014).

39. Murray, J. *Mathematical Biology*. Biomathematics (Springer Berlin Heidelberg, 2013).

40. Takeuchi, Y. *Global Dynamical Properties of Lotka-Volterra Systems* (World Scientific, 1996).

41. Guerrero, G., Langa, J. A. & Suárez, A. Attracting complex networks. In *Complex networks and dynamics*, vol. 683 of *Lecture Notes in Econom. and Math. Systems*, 309–327 (Springer, [Cham], 2016).

42. Guerrero, G., Langa, J. A. & Suárez, A. Architecture of attractor determines dynamics on mutualistic complex networks. *Nonlinear Anal. Real World Appl.* **34**, 17–40, DOI: 10.1016/j.nonrwa.2016.07.009 (2017).

43. Cottle, R., Pang, J. & Stone, R. *The Linear Complementarity Problem*. Classics in Applied Mathematics (Society for Industrial and Applied Mathematics (SIAM, 3600 Market Street, Floor 6, Philadelphia, PA 19104), 1992).

44. Murty, K. *Linear Complementarity, Linear and Non Linear Programming*. Sigma series in applied mathematics (Heldermann Verlag, 1988).

45. Cottle, R. & Balinski, M. *Complementarity and Fixed Point Problems*. Mathematical Studies (North-Holland Publishing Company, 1978).

46. Cross, G. Three types of matrix stability. *Linear Algebr. its Appl.* **20**, 253 – 263, DOI: https://doi.org/10.1016/0024-3795(78)90021-6 (1978).

47. Takeuchi, Y. & Adachi, N. The existence of globally stable equilibria of ecosystems of the generalized volterra type. *J. Math. Biol.* **10**, 401–415, DOI: 10.1007/BF00276098 (1980).

48. Ma, T. & Wang, S. Dynamic bifurcation and stability in the rayleigh-benard convection. *Commun. Math. Sci.* **2**, 159–183 (2004).

49. Ma, T. & Wang, S. *Bifurcation Theory and Applications*. Bifurcation Theory and Applications (World Scientific, 2005).

50. Rohr, R. P., Saavedra, S. & Bascompte, J. On the structural stability of mutualistic systems. *Science* **345**, DOI: 10.1126/science.1253497 (2014). https://science.sciencemag.org/content/345/6195/1253497.full.pdf.

51. Puu, T. *Attractors, Bifurcations, and Chaos: Nonlinear Phenomena in Economics* (Springer Berlin Heidelberg, 2013).
